# Multiplex Chromatin Interaction Analysis with Single-Molecule Precision

**DOI:** 10.1101/252049

**Authors:** Meizhen Zheng, Simon Zhongyuan Tian, Rahul Maurya, Byoungkoo Lee, Minji Kim, Daniel Capurso, Emaly Piecuch, Liang Gong, Jacqueline Jufen Zhu, Chee Hong Wong, Chew Yee Ngan, Ping Wang, Xiaoan Ruan, Chia-Lin Wei, Yijun Ruan

**Affiliations:** The Jackson Laboratory for Genomic Medicine, 10 Discovery Drive, Farmington, CT 06030, USA; Department of Genetics and Genome Sciences, University of Connecticut Health Center, 400 Farmington Avenue, Farmington, CT 06030, USA; College of Informatics, Huazhong Agricultural University, Wuhan, Hubei 430070, China

## Abstract

We describe a microfluidics-based strategy for genome-wide analysis of multiplex chromatin interactions with single-molecule precision. In multiplex chromatin interaction analysis (multi-ChIA), individual chromatin complexes are partitioned into droplets that contain a gel bead with unique DNA barcode, in which tethered chromatin DNA fragments are barcoded and amplified for sequencing and mapping to demarcate chromatin contacts. Thus, multi-ChIA has the unprecedented ability to uncover multiplex chromatin interactions at single-molecule level, which has been impossible using previous methods that rely on analyzing pairwise contacts via proximity ligation. We demonstrate that multiplex chromatin interactions predominantly contribute to topologically associated domains, and clusters of gene promoters and enhancers provide a fundamental topological framework for co-transcriptional regulation.

---

Genomes of higher organisms from fly to human are known to be extensively folded into chromosomal territories within the three-dimensional (3D) nuclear space^1^. Advanced long-range chromatin interaction mapping methods that combine nuclear proximity ligation and high throughput sequencing, such as Hi-C and ChIA-PET^2, 3^, have revealed frequent chromatin contacts within topological-associated domains (TADs)^4, 5^ and have implicated the formation of chromatin loops between gene promoters and enhancers as a general mechanism for transcriptional regulation^6, 7^. More specifically, studies based on RNAPII ChIA-PET have uncovered that multiple gene-coding loci and regulatory non-coding elements could be clustered together to form complex chromatin looping structures, proposing a potential topological framework for co-transcriptional regulation^6, 8^. However, current 3D genome mapping methods, Hi-C and ChIA-PET included, rely on proximity ligation to detect pairwise chromatin interactions from millions of cells. Therefore, most available 3D genome mapping data reflected only the pairwise, composite, and average views of millions of copies of chromatin molecules, and thus could not uncover the composition of multiplex chromatin interactions within each chromosome at a single-molecule resolution. Although single-cell Hi-C^9-11^ could potentially address this issue, its ability to do so may be limited by the data sparsity inherent to all single-cell genomic assays. Recent advances in microfluidics have opened new opportunities for droplet-based genomic analysis^12^, including single-cell RNA sequencing^13^ and high molecular weight genomic DNA sequencing^14, 15^, yet have not been applied to chromatin analysis. Here, we describe a droplet-based and barcode-linked sequencing strategy for multiplex chromatin interaction analysis (multi-ChIA) with single-molecule precision. In multi-ChIA, each fragmented chromatin complex is compartmentalized in a Gel bead in Emulsion (GEM) droplet that contains unique DNA oligonucleotides and reagents for linear amplification and barcoding of chromatin DNA templates. These barcoded amplicons with GEM-specific indices are pooled for high-throughput sequencing. The sequencing reads with the identical barcode are assigned to the same GEM origin, implying that they are derived from the same chromatin complex. Mapping of the DNA sequencing reads to the reference genome identifies which remote genomic loci are brought into close spatial proximity. Based on these mapped loci, multiplex chromatin interactions can be detected (Fig. 1a).

**Figure 1.**
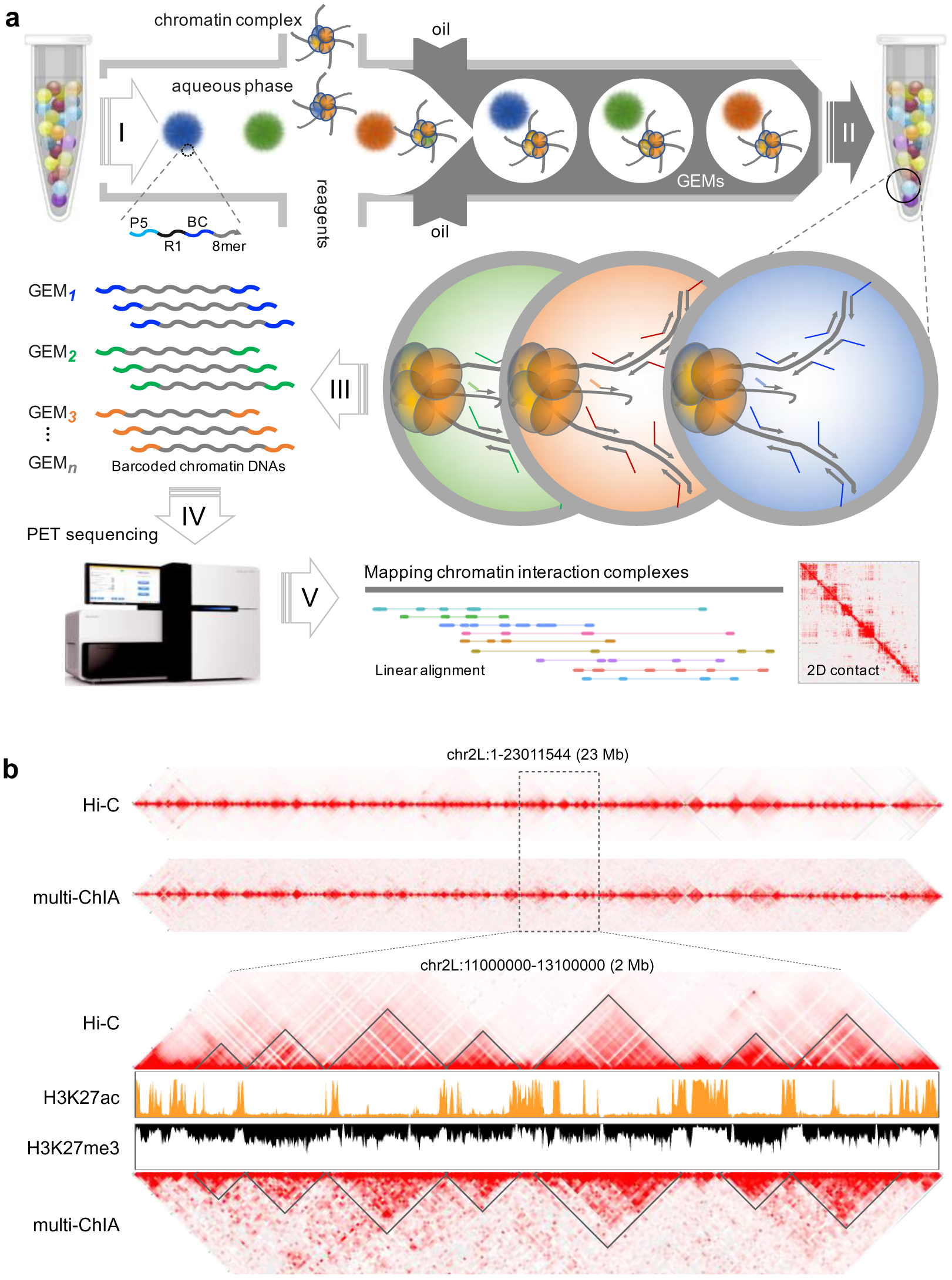
Multiplex chromatin interaction analysis. **a**, Schematic of the multi-ChIA method. (I) The microfluidics device (10x Genomics’ Chromium instrument) produces millions of Gel bead in Emulsion (GEM). Each droplet GEM is optimized to partition into one gel bead with one chromatin complex along with enzyme and reagents. Each gel bead contains millions of copies of DNA oligonucleotide as a barcode primer. P5, PCR priming site; R1, sequencing reading site; BC, 16-nucleiotide barcode; 8-mer, random 8-nucleotide for random priming to chromatin DNA templates. (II) In each GEM, chromatin DNA templates are annealed with random 8-mer for linear DNA amplification and barcoding. (III) Barcoded DNA amplicons from all GEM droplets are pooled for DNA sequencing and (V) mapping in linear alignment and 2D contact for identifying chromatin interactions. The linear alignment and 2D contact views presented here were derived from the real data of this study. **b**, Comparison of multi-ChIA data with conventional Hi-C data by pairwise 2D contact views. Top panel, whole chromosome view of chromosome 2L (chr2L). Bottom panel, a zoomed-in view of a 2 Mb segment showing comparable TADs demarcated by Hi-C data and multi-ChIA pairwise contacts. Histone marks for active domains (H3K27ac) and repressed domains (H3K27me3) are shown in the middle for reference.

To develop the multi-ChIA protocol, we used *Drosophila* Schneider 2 (S2) cells, a well-characterized model for epigenomic and 3D genome–mapping studies^5, 16^. The Chromium microfluidics system that we adopted for multi-ChIA was established for high molecular weight genomic DNA analysis^15^. To test the microfluidics system’s applicability for chromatin DNA barcoding and amplification followed by short tag sequencing, we prepared crosslinked chromatin DNA from S2 cells fragmented by restriction digestion as well as the pure DNA templates that were de-crosslinked and purified from the same chromatin DNA as a control. Digested chromatin and de-crosslinked purified DNA samples were loaded on the microfluidic chip for sample compartmentalization, GEM-specific barcoding, and amplification. Next, the DNA amplicons were analyzed by Illumina sequencing with a read length of 150 bp. We verified that the majority of the sequencing reads from pure DNA templates were of high quality, with most of the reads longer than 50 bp and many with the maximum read length (130 bp genomic plus 20 bp linker); 95.99% of the reads were uniquely mapped to *Drosophila* reference genome (dm3). By contrast, 58.65% of the sequencing reads from the chromatin DNA fragments were of high quality and mapped to the reference genome (Extended Data Fig. 1a). Most of the unmapped short reads (19–20 bp) contained only linker sequence, indicating that these GEMs might lack chromatin or the contained chromatin DNA templates were not accessible. Nonetheless, large numbers of barcoded sequencing reads generated from crosslinked chromatin material suggests that this microfluidics system is suitable for chromatin interaction analysis. We next tested whether the shorter chromatin fragments (∼300 bp) generated by restriction digestion by a 4 bp cutter (MboI) or the longer chromatin fragments (∼3000 bp) by 6 bp cutter (HindIII) are more suitable for multi-ChIA experiments. Our analysis shows that the longer fragments yield higher mappability data than shorter fragments. Similarly, chromatin prepared by sonication to fragments of around 6000 bp were also suitable for multi-ChIA analysis (Extended Data Fig. 1b). We also tested various amounts of chromatin input, and determined that 0.5 ng is the optimal loading amount for the system. When the dilution of chromatin materials was down to 0.5 pg, almost all sequencing reads contained only the linker sequence (Extended Data Fig. 1c). Finally, we assessed the probability of a GEM that might contain multiple chromatin complexes by conducting an inter-species experiment, where fly and human chromatin samples were mixed in equal amounts. If a GEM contained a mix of chromatin complexes from fly and human, then the DNA reads derived from that GEM would be mapped to both human and fly genomes (Extended Data Fig. 1d). Only 5% of the GEMs contained inter-species mixture reads from the test, indicating most GEMs that possess any chromatin material contain only one complex. In summary, this microfluidic-based system can produce high quality data for single-complex chromatin interaction analysis.

Using the multi-ChIA protocol (Methods) in this study, we generated four multi-ChIA sequencing datasets from S2 cells (Extended Data Table 1). Through the multi-ChIA data process pipeline (Extended Data Fig. 2a), we first identified the barcodes and the linked chromatin DNA reads. After mapping the reads to the reference genome, we retained only high-quality reads (≥ Q30, ≥ 50 bp), and extended them in the 3’ direction for 500 bp to better reflect the length of DNA templates subjected for sequencing analysis. Reads with the same barcode were merged into one DNA fragment if the reads overlapped. The chromatin DNA fragments tagged with the same GEM barcode were considered to be originated from the same microfluidic droplet, thus representing a single chromatin complex tethering multiple DNA fragments from different genomic loci. Given that most known chromatin-folding features are found within same chromosome territories, our further analyses on multi-ChIA data were focused on GEMs with fragments from the same chromosome (intra-chromosomal) in this study. This step has the added benefit of filtering out non-specific data potentially derived from multi-complex droplets. In total, around 3 million chromatin complexes with two or more fragments were identified (Extended Data Fig. 2f and Supplementary Table 1). More than a half of the them (1,594,169) contained three or more fragments, thus provided a large dataset to study potential multiplex chromatin interactions that might involve three or more contacting loci simultaneously, which could not be addressed by the pairwise-based Hi-C data. Surprisingly, many of the multi-ChIA complexes (38,293) include ten and up to hundreds of fragments (Supplementary Table 1), indicating extensive high complexity of chromatin interactions.

With the exception of its unique ability to resolve the multiplex nature of chromatin complexes, multi-ChIA should essentially detect the same spectrum of chromatin contacts as Hi-C. We thus compared the multi-ChIA data with the Hi-C data derived from the same S2 cells^17, 18^ to validate the new method. The chromatin complexes with multiple DNA fragments in multi-ChIA data should reflect multiple, simultaneous chromatin contacts. In contrast, because Hi-C relies on proximity ligation between two DNA fragments, each fragment in a Hi-C experiment can only be associated with one other fragment. Thus, the ligation-based pairwise contacts in a Hi-C experiment, which uses millions of cells, are an accumulation from random sampling of all possible proximity ligation products among millions of chromosome copies. To directly compare multi-ChIA with Hi-C data, we devised a random-sampling algorithm with multiple iterations to best mimic the pairwise Hi-C-like output from the multi-ChIA data (Methods) and visualized the adjusted multi-ChIA data in 2D contact maps (Extended Data Fig. 3). As shown, the multi-ChIA data captured chromatin structural features that largely resemble those in Hi-C data (Fig. 1b and Extended Data Fig. 4). Notably, most of the distinctive topologically associated domains (TADs)^18^ mapped by both Hi-C and multi-ChIA were closely associated with repressed chromatin domains (Fig. 1b).

Unlike Hi-C, a major advantage of multi-ChIA is to capture simultaneous contacts between multiple genomic loci, providing the numbers of DNA fragments per complex from two up to hundred fragments. This unique feature allows us to characterize the contribution of chromatin complexes with varying numbers of interacting elements to the chromatin structures viewed in 2D contact displays. We decomposed the multi-ChIA data according to the number of fragments in each complex (F=2, complexes with 2 fragments; F=3, with 3 fragments; and so forth), and plotted the data of each class for 2D contact analysis. Intriguingly, most of the contacts from low-fragment-containing complexes (F=2–5) were either in short-distance along the 2D contact axis, or long-distance randomly scattered, whereas complexes with higher number fragments tend to cluster and provide the framework of the chromatin topological structures (Fig. 2a and Extended Data Fig. 5). Statistical analysis by empirical cumulative distribution function (ECDF) showed that the pairwise contacts of the low-fragment-containing complex data were mostly at extremely long distance and deviated more from the reference ECDF curve of Hi-C data (Fig. 2b), further suggesting that the high-fragment-containing complex data were of sufficiently high quality for multiplex chromatin interaction analysis.

**Figure 2.**
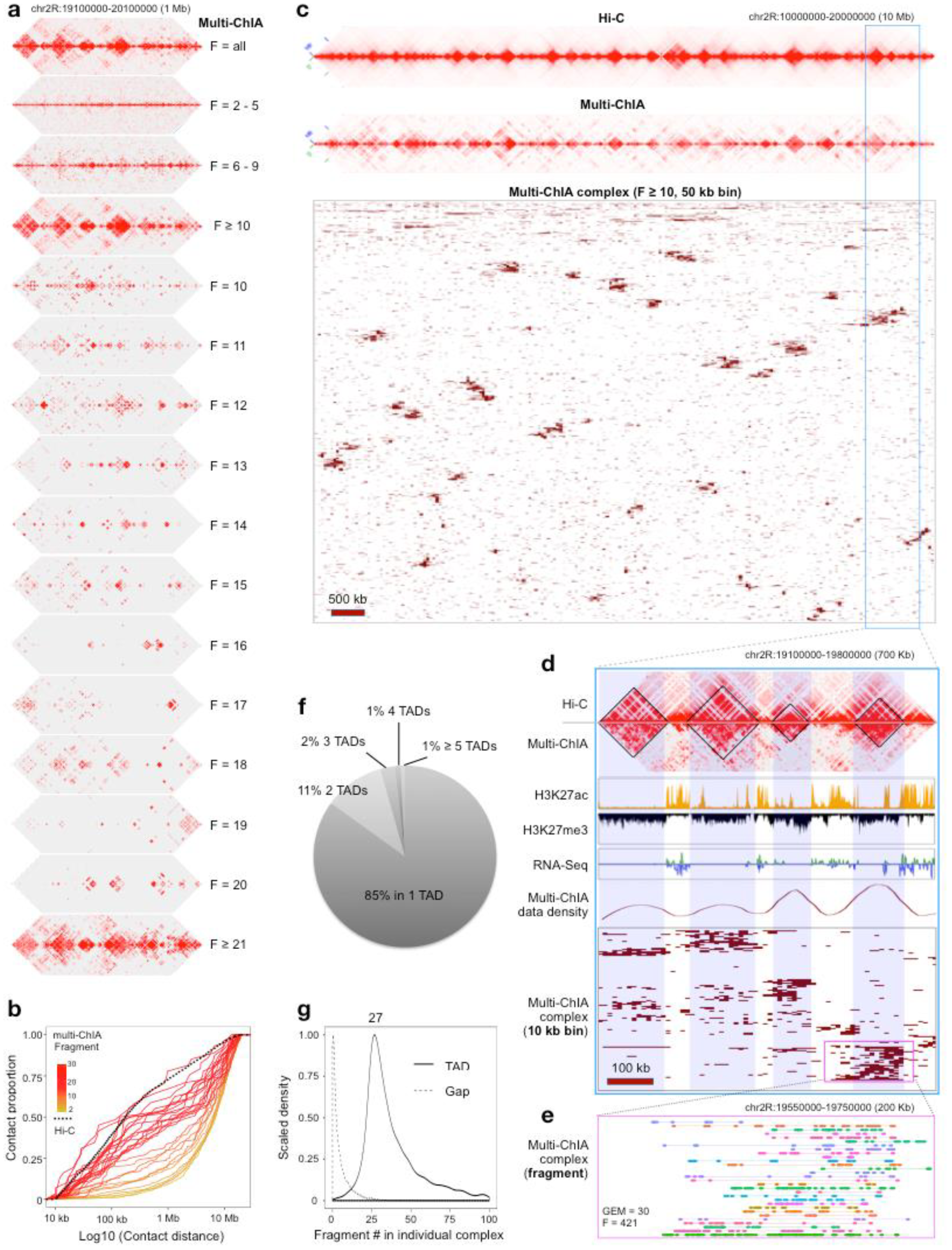
Most multiplex chromatin-interaction complexes form within TADs. **a,** Decomposition of multi-ChIA data based on the number of DNA fragments in each chromatin complex. Shown are 2D contact heatmaps of a 1 Mb segment of chr2R for all data (F = all), complexes with only two to five fragments (F = 2–5), six to nine fragments (F = 6–9), or ten and more fragments (F ≥ 10). These data are further broken down into complexes with specific numbers of fragments, ranging from F = 10 to F ≥ 21. **b,** Empirical cumulative distribution function (ECDF) of the pairwise chromatin contact distance in log scale. The solid curves denote multi-ChIA data with color intensity (yellow to red) corresponding to different numbers of fragments per complex from low to high. The black dotted curve represents Hi-C data as a reference. **c,** Multi-ChIA data in a 10 Mb segment of chr2R. The top panel displays the comparable profiles of pairwise contacts between Hi-C and multi-ChIA data. The bottom panel shows alignments of chromatin complexes, each with more than 10 fragments (F ≥ 10). Chromatin fragments in each complex were binned with 50 kb resolution, and were reordered by the hierarchical clustering of the bins, showing the enrichment of chromatin complexes in TAD structures. **d,** A zoomed-in view of Hi-C and multi-ChIA data from (b) shows a 700 kb segment of four TADs along with histone and transcription profiles. Linear alignment of chromatin complex fragments with 10 kb bins and hierarchical clustering highlights enrichment of multi-ChIA data in TAD domains (middle). **e,** Further zoom-in (200 kb) displays detailed alignment of the 421 fragments as assorted color bars connected in 30 chromatin complexes. **f,** A pie chart shows the proportion of chromatin complexes with multiple fragments in TAD domains. 85% complexes have all fragments located within a single TAD, and the rest have fragments over multiple TADs. **g,** A density plot shows that the numbers of fragments per complex in those complexes mapped to TADs (solid line) was much higher than the number of fragments per complex in those complexes mapped to the gaps between TAD boundary regions (dotted line).

We then focused on multi-ChIA complex data with high numbers of simultaneous contacts (F≥10, n=38,293) for further analysis (Supplementary Table 1). By binning contact data at different resolutions (50 kb, 10 kb and 5 kb) and by performing hierarchical-clustering algorithm, we could also directly view individual chromatin complex data in linear alignment across the genome (Methods, Extended Data Fig. 6). For example, in a 10 Mb segment of chr2R, there were 2,088 chromatin complexes identified by multi-ChIA. At a 50 kb bin size resolution, many clusters of chromatin complexes were observed to be highly correlated with TAD structures defined by both multi-ChIA data and Hi-C data (Fig. 2c). A 700 kb region with 10 kb bins resolution showed more detailed patterns of multi-ChIA complexes in tighter association with TAD structures in repressed chromatin domains as defined by histone modification markers and transcriptional profiles (Fig. 2d). With further zooming, individual fragments captured within single chromatin complexes could be visualized. For instance, within a 200 kb region in chr2R that includes a TAD, a selected set of 421 fragments were visualized as assorted color bars for their relative length and were connected in 30 chromatin complexes (Fig. 2e). Overall, most of the multi-ChIA complex data (85%) fall within individual TADs, and some cover two or more TAD structures (Fig. 2f). By contrast, relatively fewer chromatin fragments fall within the gap regions between TAD boundaries (Fig. 2g). Altogether, our analyses provided evidence to support that most of chromatin interactions within TADs involve simultaneous contacts among multiple loci, forming complex structures of individual chromatin fibers.

To investigate chromatin interactions involved in transcriptional regulation in *Drosophila* S2 cells, we performed RNAPII ChIA-PET experiments based on proximity ligation and paired-end-tag sequencing (Methods). The ChIA-PET data mapped abundant chromatin interaction loops mediated by RNAPII (Extended Data Fig. 7). Many RNAPII loops are interconnected in daisy-chain structures in transcription active regions, and are defined as RNAPII-associated interaction domains (RAIDs) (Fig. 3a and Extended Data Fig. 7d). From the RNAPII ChIA-PET data, we identified 476 RAIDs in S2 cells (Supplementary Table 3), most of which are composed of multiple interacting loci of transcriptional elements such as promoters and enhancers (Fig. 3a and Extended Data Fig. 7d). This finding suggests a possible mechanism for orchestration of transcription of genes involved in the domain, as observed previously in human^6, 8^ and mouse genomes^19^. We also characterized the S2 cell genome based on H3K27ac (transcriptionally active) and H3K27me3 (transcriptionally repressive) ChIP-seq data. Along with RNA-Seq data, we demarcated the S2 epigenome landscape as active and inactive domains (Methods, Supplementary Table 3), and confirmed that most RAIDs overlap with the active domains (Fig. 3a). In addition, there are relatively lower frequent and longer chromatin contacts between RAIDs (Fig. 3a and Extended Data Fig. 7), hinting further complexity of higher order chromatin organization involved in transcriptional regulation. Regardless, ChIA-PET data could not alone determine whether multiple gene promoters and enhancers on same chromosome were indeed in simultaneous contact at the single molecule level, or whether we were capturing different, transient pairwise contacts averaged across millions of chromosomes in a ChIA-PET experiment.

**Figure 3.**
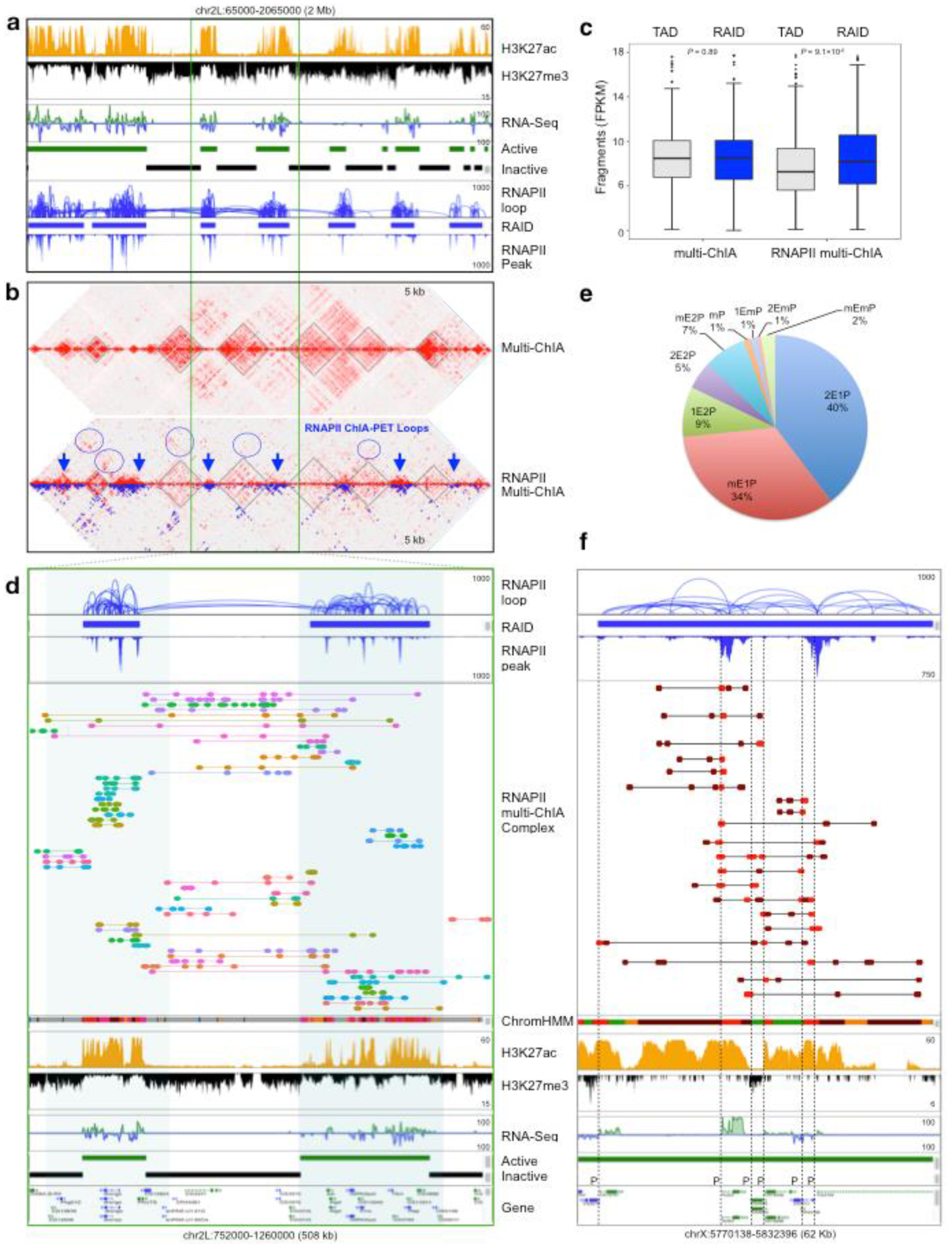
RNAPII multi-ChIA data demonstrate complex chromatin interactions involving multiple gene promoters and enhancers. **a,** The epigenomic landscape of active/inactive domains and RNAPII ChIA-PET data defines RAIDs and inter-RAIDs connectivity, as shown in a 2 Mb segment of chr2L. **b,** Comparison of standard multi-ChIA and RNAPII enriched multi-ChIA data using 2D heatmap for pairwise contacts. TADs defined by both data are outlined in black. RNAPII-mediated chromatin loops (blue dots) identified by ChIA-PET are superimposed on the RNAPII multi-ChIA data. RNAPII enriched regions in RNAPII multi-ChIA data are indicated by blue arrows. Inter-RAIDs contacts in RNAPII multi-ChIA data are also circled in blue. **c,** A boxplot of chromatin fragment represents in defined TADs and RAIDs in both non-enriched multi-ChIA and RNAPII enriched multi-ChIA data. Boxes depict median and interquartile range, and *p*-values are from the Wilcoxon Rank-Sum Test. **d,** A zoomed-in view of (**a**) shows a browser display of two RAIDs connected by pairwise RNAPII loops from the ChIA-PET data and chromatin interaction complexes with multiple contacting fragments identified by RNAPII multi-ChIA data. Tracks of chromatin state (ChromHMM: red for promoter; dark for enhancer; grey for inactive region), histone marks (ChIP-Seq for H3K27ac and H3K27me3), transcription activity (RNA-seq), and gene model track are provided for genomic and epigenomic references. **e,** A pie chart for chromatin complexes with different composition of promoters and enhancers involved multiplex interactions detected by RNAPII multi-ChIA. Of the 11,695 chromatin complexes with 3 or more fragments (F≥3) overlapping with a RAID, most of the complexes (74%) mapped with one promoter (P) and 2 or more enhancers (E). The rest were consisting of 2 or more promoters with various numbers of enhancers. 2E1P represents complexes contained 2 enhancers and 1 promoters, mE1P as multiple enhancers and 1 promoter, and so forth. **f,** An example of a browser view showing a RAID with abundant RNAPII loops by ChIA-PET around *Act5C* and 6 other genes. In this region, RNAPII multi-ChIA data detected 20 chromatin complexes containing multiple regulatory elements simultaneously. All of complexes included at least one promoter (proximal to transcription start site, TSS) and multiple possible enhancers (distal to TSS), 13 of them connected 1 promoter, 5 included 2 promoters, and 2 included 3 promoters. *Act5C* is the most actively expressed gene in this block, and the *Act5C* promoter were directly covered by 10 of the complexes. Other genes in this region were less expressed, and their promoters had fewer complexes covered. The red bar depict promoter and the brown bar denote enhancer. The corresponding TSSs of genes are indicated by dotted vertical lines.

To address these questions, we added a chromatin immunoprecipitation (ChIP) step against RNAPII to our multi-ChIA protocol. We performed RNAPII-enriched multi-ChIA experiments from S2 cells (Methods), and generated 2 RNAPII multi-ChIA sequencing data (Extended Data Table 1). Following the same multi-ChIA data process pipeline (Extended Data Fig. 2a), we identified 433,932 complexes with multiple chromatin fragments (Extended Data Fig. 8a-b, Supplementary Table 2). Using pairwise 2D chromatin contacts profiling for comparison, similar to our previous multi-ChIA vs. Hi-C comparison, the RNAPII multi-ChIA data and RNAPII ChIA-PET data exhibited very similar chromatin contact patterns (Extended Data Fig. 8c-d), indicating that both methods captured the same spectrum of chromatin interactions associated with RNAPII in S2 cells.

When comparing the RNAPII multi-ChIA data (with RNAPII enrichment) and the standard multi-ChIA data (without RNAPII enrichment), it is striking that the RNAPII multi-ChIA data exhibited considerable reductions of chromatin contact signals in TAD regions^18^ and inactive domains, whereas substantial enrichment of chromatin contact signals for RNAPII loops in the RAID regions and active domains (Fig. 3b–c and Extended Data Fig. 10). In addition, RNAPII multi-ChIA data also showed inter-RAID contacts across the intervening TAD structures (Fig. 3b), recapitulating the patterns observed in the RNAPII ChIA-PET data (Fig. 3a). These observations provide validation that the multi-ChIA experimental system successfully reflects the properties of input chromatin samples (enriched or not), and that multi-ChIA can be applied to investigate multiplex interactions among regulatory elements associated with transcriptional activity.

As with the non-enriched multi-ChIA data, we also conducted decomposition analysis for the RNAPII multi-ChIA complex data based on fragment-containing multiplexity. Similar to the multi-ChIA data (Fig. 2a-b and Extended Data Fig. 5), the RNAPII multi-ChIA complexes with few fragments lacked distinctive 2D contacts, in contrast to the high-fragment-containing complexes that contributed to topological features (Extended Data Fig. 9a-b). Statistical analysis by empirical cumulative distribution function (ECDF) of pairwise contact distances between chromatin fragments showed that the high-fragment-containing complexes matched tightly with the reference ECDF curve of the RNAPII ChIA-PET data, whereas the low-fragment-containing complex data showed significant deviation (Extended Data Fig. 9c), providing another indication that the high-fragment-containing complex data likely represent genuine multiplex chromatin structures.

We further analyzed the RNAPII multi-ChIA data in relation to RAIDs identified by RNAPII ChIA-PET data. It was observed that a high percentage of chromatin fragments in RNAPII multi-ChIA data were overlapped within individual RAIDs, and there were also some that connected between RAIDs (Extended Data Fig. 11 and Fig. 3d). More specifically, we have detected large numbers of chromatin interaction complexes in RNAPII multi-ChIA data that simultaneously connected multiple transcriptional elements such as gene promoters and potential enhancers in RAIDs, more than 3000 such complexes included at least two promoters (Methods, Fig. 3e and Supplementary Table 4). For example, at the locus of *Act5C* (encoding actin-5C protein for cytoskeletal structure)^20^ and other seven genes (Fig. 3f), we identified twenty chromatin complexes from the RNAPII multi-ChIA data. All of them simultaneously connected to multiple transcriptional elements including promoters and enhancers, and eight of them included two or three promoters. Notably, *Act5C* is the most actively expressed gene in the block, and the *Act5C* promoter were covered by ten of the individual complexes and also displayed direct connectivity with three other gene promoters, whereas the other genes here were less expressed and with fewer complexes detected. In addition to inter-genic chromatin contacts, we also observed abundant intra-genic contacts in RNAPII multi-ChIA data, particularly for large genes such as *pum* and *luna* over 100 kb length^21^, revealing potential topological details of chromatin templates during transcription (Extended Data Fig. 12c-d). More examples are presented in Supplementary Information (Extended Data Fig. 12 and Supplementary Table 4).

In summary, we have demonstrated that multi-ChIA is an effective method for investigating multiplex chromatin interactions at the single-molecule level, while previous methods like Hi-C and ChIA-PET could not. Through our analyses in *Drosophila* cells, we have shown that most TADs are associated with repressed domains and composed of simultaneous multiplex chromatin interactions on the same chromatin string. More importantly, we have provided convincing evidence in RNAPII multi-ChIA data for simultaneous intergenic chromatin interactions involving multiple gene promoters and enhancers on single chromosomes. These findings support a topological mechanism of co-transcriptional regulation, in which multiple gene promoters and regulatory elements are brought together in simultaneous contact for active transcription. Finally, the multi-ChIA protocol is exceptionally simple and robust. Fragmented chromatin samples as prepared as for Hi-C or ChIA-PET, but without the need of proximity ligation, can be directly processed in an automated microfluidics device for gel bead emulsion and droplet-specific chromatin DNA barcoding and amplification followed by standard Illumina sequencing. The protocol requires only 5 × 10^3^ cells for a multi-ChIA experiment or approximately 6 × 10^4^ cells for a ChIP-enriched multi-ChIA experiment under optimized conditions. We anticipate that multi-ChIA will be rapidly adapted for a wide range of biomedical questions and applications.

## Methods

Methods, including statements of data availability and any associated accession codes and references, are in online version of Methods.

## Acknowledgments

This study is supported by a Jackson Laboratory Director’s Innovation Fund (DIF19000-18-02). YR and CLW are funded by 4DN (U54 DK107967) and ENCODE (UM1 HG009409) consortia. YR is also funded by Human Frontier Science Program (RGP0039/2017), and supported by the Florine Roux Endowment.

## Author contribution

M.Z., Y.R. and C.L.W. conceptualized the multi-ChIA method. M.Z. implemented the multi-ChIA protocol, designed the studies, and conducted experiments with assistant from R.M. on performing 10x Genomic’s Chromium protocol. R.M. and C.Y.N. coordinated multi-ChIA library sequencing. P.W., E.P. and X.R. contributed RNAPII ChIA-PET data. S.Z.T. developed the computational pipeline for multi-ChIA data processing and analysis with helps from C.H.W. and D.C., and M.K. and B.L. developed the algorithms for simulating Hi-C-like contacts from multi-ChIA data. S.Z.T., D.C., M.K. and B.L. performed various statistical analyses. S.Z.T. and D.C. developed the linear visualization program for multi-ChIA data. B.L. wrote a script to convert multi-ChIA data for 2D contact visualization using Juicebox. M.Z. and Y.R. wrote the manuscript with input from M.K., S.Z.T., B.L., and D.C. All co-authors read and approved the manuscript.

## Online version of Methods

### I. EXPERIMENTAL METHODS

#### Cell culture

*Drosophila* Schneider 2 (S2) cells (derived from a primary culture of late stage 20-24 hours old *Drosophila* melanogaster embryos) were cultured in Express Five^®^ SFM (ThermoFisher Scientific) with 1:100 L-Glutamine (ThermoFisher Scientific) at 27 °C. Human GM12878 cells (a B-lymphoblastoid cell line produced from the blood of a female donor with northern and western European ancestry by EBV transformation) were maintained in RPMI 1640 supplied with 15% fetal bovine serum at 37 °C and ambient 5% CO2 as described by the Coriell Institute of Medical Research. Cells at exponential growth phase were harvested for chromatin preparation.

#### Chromatin sample preparation for multi-ChIA

The overall chromatin sample preparation for multi-ChIA is very similar to sample preparation for Hi-C^22^, except no proximity ligation. In brief, 10 million cells were crosslinked with 1% formaldehyde at room temperature for 10 min, and quenched with 0.125 M Glycine (Promega) for 5 min, then washed with DPBS twice. The crosslinked cells can be stored at -80 °C for later use or immediately proceed for cell/nuclei lysis. The crosslinked cells were suspended in 500 μl of cell lysis buffer (10 mM Tris-HCl pH 7.0, 10 mM Tris-HCl pH 8.0, 10 mM NaCl, 0.2% NP40, 1× Protease Inhibitor cocktail, Roche) and incubated at 4 °C for 30 min with rotation. Then, the nuclei were isolated by centrifugation at 4 °C for 5 min at 2,500 relative centrifugal force (rcf). The nuclei pellet was suspended in 100 μl of 0.5% SDS and incubated at 62 °C for 5 min to permeabilize the nuclear membrane. After that, 285 μl of nuclease-free water and 25 μl of 20% triton X-100 were added for further incubation at 37 °C for 15 min to neutralize SDS. The permeabilized nuclei were then ready for in situ chromatin digestion. For digestion by 4 bp cutter MboI (catæ R0147L, NEB), 60 μl of NEB Buffer 2 was added and mixed well, then 55 μl of nuclease-free water and 75 μl of MboI (5 U/μl) were added, whereas for 6 bp cuter digestion HindIII (catæ R0104L, NEB), 80 μl of nuclease-free water and 50 μl of HindIII (20 U/μl) were added to set up the reactions. The incubations were taking place at 37 °C overnight with constant agitation. The nuclei with digested chromatin materials were sheared by sonication with 1× Protease Inhibitor cocktail to release the chromatin fragments. The DNA size range of the chromatin fragments usually was 300-6000 bp, depending on restriction digestion. The final fragmented chromatin sample was then preceded for multi-ChIA library construction. If not immediately used, the chromatin sample should be stored at 4 °C, and no longer than a week.

#### Chromatin sample preparation for RNAPII multi-ChIA

The overall chromatin sample preparation for RNAPII enriched multi-ChIA follows the ChIA-PET protocol^23^, except no proximity ligation. In brief, 10 million cells were dual-crosslinked with 1.5 mM EGS for 40 min followed by 1% formaldehyde reaction for 20 min, and then quenched with 0.125 M Glycine (Promega) for 10 min, and washed twice with DPBS. The crosslinked cells can be frozen and stored at -80 °C for later use. After cell and nuclei lysis, the crosslinked chromatin material was fragmented by sonication to the size range of 6 kb. The fragmented chromatin sample was incubated with 20 μl of anti-RNAPII monoclonal antibody (8WG16, catæ 920102, Biolegend, San Diego, CA) bound on Dynabeads^™^ Protein G beads (catæ 10009D, ThermoFisher Scientific) at 4 °C overnight with rotation. RNAPII-enriched chromatin was released from Protein G beads by incubating with EB Buffer containing 1% SDS at 37 °C for 30 min with constant agitation. The elution supernatant was passed through Ultra Centrifugal Filter (catæ UFC510024, Millipore) to remove the remaining SDS. The chromatin preparation now is ready for multi-ChIA library construction, or stored at 4 °C for later use.

#### Multi-ChIA library construction and sequencing

Fragmented chromatin sample was mixed with 50 μg/ml of BSA (catæ B9000S, NEB) to prevent chromatin aggregation. An aliquot of chromatin sample was taken to estimate the quantity of chromatin DNA, and the chromatin sample was adjusted to the concentration of 0.5 ng/μl. We applied the GemCode Technology (10x Genomics, Pleasant, CA) for partitioning of individual chromatin complexes into droplets for chromatin DNA barcoding and amplification. In brief, the chromatin sample was mixed with Sample Master Mix from Chromium Genome v2 Library Kit & Gel Bead Kit (catæ PN-120258), and input 0.5 ng chromatin mix was loaded into the Chromium Genome Chip (catæ PN-120257), which was placed in Chromium Controller to generate Gel bead in Emulsion (GEMs) droplets each containing unique barcoded DNA primers, chromatin DNA fragments, and enzymatic reagents. The GEMs were subjected to a 30 °C isothermal incubation for overnight for amplification and barcoding of the chromatin DNA templates. After that, the barcoded DNA amplicons were released from all GEMs. Purified DNA fragments were subjected for end repair, A-tailing, adaptor ligation and Sample Index PCR using reagents from Chromium Multiplex Kit (catæ PN-120262) following the manufacturer’s instructions. The PCR products were size-selected in the range of 600 bp, and the final multi-ChIA library was analyzed by sequencing (2×150 bp) using MiSeq (Illumina, San Diego, CA). The R1 reads that include GEMcode and genomic sequences were used for further analysis in this study.

#### RNAPII ChIA-PET library preparation and sequencing

The RNAPII ChIA-PET libraries were performed as previously described^23^. Briefly, dual-crosslinked *Drosophila S2* cells were lysed followed by sonication to fragment the chromatin material into the range of 6 kb in length. The fragmented chromatin sample was subjected for ChIP enrichment by anti-RNAPII monoclonal antibody (8WG16, catæ 920102, Biolegend, San Diego, CA). The RNAPII-enriched chromatin material was then subjected for ChIA-PET library construction, followed by paired-end-tag sequencing (2× 150 bp) using Illumina MiSeq instruments. We also applied an improved ChIA-PET protocol, called in situ ChIA-PET, to generate RNAPII ChIA-PET libraries for S2 cells. In brief, 10 million cells were subjected for dual-crosslinking^2^ followed by cell lysis in 1 ml of 0.1% FA Buffer (50 mM HEPES, 150 mM NaCl, 1 mM EDTA, 1% triton X-100, 0.1% sodium deoxycholate and 0.1% SDS, 1× Protease Inhibitor cocktail) at 4 °C for 20 min with rotation. Nuclei were isolated by centrifugation (5,000 rcf) at 4 °C for 10 min. The nuclei pellet was then re-suspended in 500 μl of 0.1% FA buffer with 0.55% SDS, and the nuclei were permeabilized at room temperature for 10 min with further incubation at 62 °C for additional 10 min and at 37 °C for 10 min. The permeabilized nuclei were pelleted by centrifugation (5,000 rcf) for 10 min, and washed with 1× NEB buffer 2.1 for one time, then re-suspended in 500 μl of 1× NEB Buffer 2.1 with 50 μl of AluI (catæ R0137L, NEB) for in situ digestion at 37 °C for overnight. AluI was inactivated with 10 μl of 0.5 M EDTA at 37 °C for 10 min, and nuclei was pelleted by centrifugation (5,000 rcf) for 10 min, then subjected for in situ end-repair, A-tailing, and proximity ligation with bridge linker^23^. The nuclei with in situ proximity ligation were lysed followed by sonication to release the chromatin material. The fragmented chromatin preparation was incubated with anti-RNAPII antibody (8WG16, catæ 920102, Biolegend, San Diego, CA) for immunoprecipitation. The RNAPII enriched chromatin material was subjected for ChIA-PET library construction and DNA sequencing following the ChIA-PET protocol^23^.

### II. COMPUTATIONAL DATA PROCESSING

#### Multi-ChIA data processing pipeline

The multi-ChIA data were generated by 2x150 bp sequencing. For simplicity, only R1 reads that contain GEM barcodes (GEMcodes) and genomic sequences were used for further analysis in this study. The R2 reads from the other end of the multi-ChIA library DNA templates contain only genomic sequences will be included for future analysis. To analyze the multi-ChIA sequencing data, we developed a data processing pipeline (Extended Data Fig. 2a). It uses modules from the Chromium software suite (10x Genomics) for aligning sequencing reads and identifying GEMcodes. In addition, it uses custom scripts specifically developed for multi-ChIA data, for merging overlapping reads originated from the same GEMs into chromatin fragments, and for characterizing GEMs by counting the number of fragments within them. GEMs with multiple fragments represent chromatin interaction complexes.

**(1) Alignment of sequencing data and identification of GEMcodes**. FASTQ files were aligned to the *Drosophila* reference genome (dm3) using the 10x longranger wgs pipeline (v2.1.5, https://support.10xgenomics.com/genome-exome/software/pipelines/latest/using/wgs) with default parameters to generate BAM files and simultaneously identify GEMcodes.

**(2) Renaming read IDs by prepending them with library IDs and GEMcodes**. The pysam module (v0.7.5, https://github.com/pysam-developers/pysam) of python (v2.7.9) was used to access the BAM files and extract GEMcodes from the tag field “BX”.Then, each read was given a new ID, which is comprised of the library ID, the GEMcode, and the original read ID. An example of a new read ID in a GEMcoded-BAM file is:

“SHG0021N-101294002-TACTTGTTCTTGTATC-NS500460:311:HGK53BGX3:4:21605:26065:15070”,

where “SHG0021N” is the library ID, “101294002” is GEMcode ID, “TACTTGTTCTTGTATC” is the GEMcode sequence, and “NS500460:311:HGK53BGX3:4:21605:26065:15070” is the original read ID. This re-naming allows the GEMcode to be propagated forward, when the BAM files are converted to BED files for downstream analysis.

**(3) Quality-filtering of the aligned reads**. Only uniquely mapped R1 reads with MAPQ ≥ 30 and read length ≥ 50 bp were retained for downstream analyses. These reads were converted to BED format using bedtools (v2.17.0)^24^.

**(4) Merging the sequencing reads into chromatin fragments**. We extended each high-quality read by 500 bp from its 3’ end, reflecting the DNA template size that was used for sequencing (∼ 300 -600 bp). Considering that in the experimental protocol, the chromatin fragments were randomly primed for linear amplification, the same chromatin fragment could be represented by multiple different but overlapping sequencing reads. Therefore, reads with the same GEMcode and whose extended genomic coordinates overlapped were merged to represent a chromatin fragment. Merging was performed using pybedtools (v0.7.10)^25^.

**(5) Characterization of chromatin interaction complexes**. Based on the merged chromatin fragments and their GEMcodes, individual GEMs could then be categorized based on the number of fragments they contained. A GEM with one chromatin fragment (F=1) was considered a singleton GEM, whereas a GEM with two or more fragments (F≥2) was considered a multiplex GEM, representing a chromatin interaction complex with multiple fragments. Based on the genomic coordinates of the chromatin fragments, multiplex GEMs were further classified as intra-chromosomal GEMs if all fragments within a GEM were mapped to the same chromosome or inter-chromosomal GEMs if the fragments within a GEM mapped to different chromosomes. In this study, we considered only intra-chromosomal GEMs for investigations of chromatin interaction complexes in same chromosomes. In addition, inter-chromosomal GEMs were further filtered for their intra-chromosomal components. For instance, a given GEM possesses 5 chromatin fragments, in which 2 were mapped to one chromosome and the other 3 were to another chromosome. This GEM would be split into two intra-chromosomal subsets. The final data set for downstream analyses includes all intra-chromosomal interaction complexes.

#### RNAPII ChIA-PET data analysis

RNAPII ChIA-PET data and in situ RNAPII ChIA-PET data derived from *Drosophila* S2 cells were processed following the established ChIA-PET pipeline^23^. Briefly, linker sequences were trimmed off and paired-end tags (PETs) were aligned to the *Drosophila* reference genome (dm3). Uniquely mapped and non-redundant PET reads were classified as self-ligation PETs or inter-ligation PETs. Inter-ligation PETs were further clustered together if both tags (with a 500 bp extension) overlapped. Clusters with PET counts of 2 or more (PET≥2) were considered as candidates for further analysis of chromatin interaction loops. Chromatin loops with higher PET counts reflect frequent contacts, and are considered high confidence data.

### III. VISUALIZATION OF MULTI-CHIA DATA

The merged chromatin fragments in each of the chromatin complexes and their contact profiles were visualized in 2D contact maps and in linear connectivity.

(1) **2D heat-map visualization of multi-ChIA data.**The pairwise interactions in BEDPE format were used as input for generating a contact map file in .hic format using Juicer tools (version 1.7.5)^26^. Subsequently, visualization was performed using Juicebox (version 1.6.2)^27^. Various resolutions of bin size from 2.5 Mb to 1 kb were used to visualize the data in the 6 chromosomes of the *Drosophila* reference genome (dm3).

(2) **Linear connectivity visualization of multi-ChIA data.**To visualize multiplex chromatin interaction complexes in genome-based browser, we developed a custom tool in R, which displays the chromatin fragments in each GEM unit along the linear genome browser after ordering the GEMs by similarity. Briefly, the genomic segment of interest in mega-base range is binned (default: 100 bins), and a Boolean matrix is constructed, where the bins are the columns and each row is a GEM. For each GEM, it is recorded whether or not each bin contains a fragment. Then, the GEMs are ordered based on similarity via hierarchical clustering (default: complete linkage). To visualize individual GEMs in genomic segments under mega-base size, specific chromatin fragments are plotted using their original fragment coordinates in assorted color bars along the genome, and the fragments belonging to the same GEMs are grouped by connecting lines (visualization examples shown in Fig. 2-3, Extended Data Fig. 6, 11 and 12).

### IV. STATISTICAL ANALYSIS

All statistical analyses in this study were performed using the R statistical package (http://www.r-project.org). Further details are described below.

#### Summary statistics of multi-ChIA data

The multi-ChIA sequencing reads have a maximum length of 130 bp (Extended Data Fig. 2c). High quality sequencing reads (MAPQ ≥30, read length ≥ 50 bp) were extended for 500 bp from the 3’ ends, and merged into chromatin fragments (as described above). The average fragment length is ∼ 630 bp (Extended Data Fig. 2d). Fragments sharing the same origin of GEMcode were group as a GEM unit, reflecting the chromatin composition. In total, 6,068,419 GEMs were identified in the multi-ChIA data. More than half of the GEMs (3,144,087) contain multiple chromatin fragments (F≥2), representing more than 3 million multiplex chromatin complexes (Extended Data Fig. 2e-f). Further details on the genomic coordinates of multiplex GEMs with 10 or more fragments (F≥10) are provided in Supplementary Table 1.

#### Summary statistics of RNAPII multi-ChIA sequencing data

For RNAPII multi-ChIA data, the maximum read length and average length of the merged fragments are similar to those of multi-ChIA. Of the total GEMs (2,831,907), 85% (2,397,975) were singleton GEM (F=1, containing one fragment), and the rest were multiplex GEMs containing 2 and more fragments (Extended Data Fig. 8a-b). Further details on the genomic coordinates of high multiplex GEMs (12,930) with six or more fragments (F≥6) are provided in Supplementary Table 2.

#### Comparison of multi-ChIA and Hi-C data

In theory, the multi-ChIA and Hi-C protocols should capture comparable chromatin interactions. To support our hypothesis, we compared 2D contact heat maps of the two datasets. However, there are fundamental differences between the two datasets that needs to be considered. The output of a multi-ChIA experiment is a collection of chromatin complexes, each with multiple fragments (independent of proximity ligation), whereas the Hi-C protocol generates only pairwise interactions by proximity ligation between two ends of the same (self-ligation) or different (inter-ligation) DNA fragments. For instance, a chromatin complex with 10 fragments identified by multi-ChIA implies that these 10 loci were all in close contact. However, it would not be possible for a Hi-C experiment to capture all possible 45 pairwise interactions from this complex with 10 fragments within the same cell; by the nature of the experimental technique, the ligated DNA ends are no longer available to be ligated with other proximal DNA fragments. Considering that a Hi-C experiment is a random sampling process that produces a subset of all possible proximity ligation events from millions of cells, we conceived an algorithm to best mimic the Hi-C pairwise output from the multi-ChIA data, as detailed below.

We convert multi-ChIA complex data into pairwise Hi-C-like data based on an algorithm that simulates <u>P</u>roximity <u>L</u>igation *<u>I</u>n <u>S</u>ilico* by <u>R</u>andom <u>S</u>ampling (PLISRS), a specific program developed for this purpose. In PLISRS, we consider each chromatin complex is a set of ***n***fragments and each fragment has two endpoints available to be ligated. Thus, we can represent a complex as a set of 2*n* endpoints {**r*_*1*_, *r*_*1*_, *r*_*2*_, *r*_*2*_,…,*r*_*n*_, *r*_*n*_*} and randomly sample 2 endpoints from the set without replacement (‘random’ module in python v3.6.3). Note that it is possible to choose two endpoints from the same fragment (self-ligation), i.e., *r*_*t*_ and *r*_*t*_, for *i* in {*1,2,…,n*}. We iterate the process *k* times for all complexes and the sampled pairs are de-duplicated. The rationale behind de-duplication is to follow a typical Hi-C data processing pipeline, where we take only one representative read if multiple reads are mapped to the same genomic location. Iteration number *k* can be a variable parameter, but we chose 5 because we add 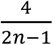 of inter-ligations in every iteration, and 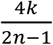 > 1 for *n ≤* 10 when *k =* 5. In loose, we would have sampled all pairs for *n ≤* 10, which account for the majority of the complexes. When *n >* 10, we don’t necessarily sample all possible pairs 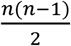 (which would be equivalent to the original multi-ChIA data), but sample enough pairs to simulate Hi-C-like data, with an assumption that Hi-C is an incomplete sampling from all probability of millions of cells.

To characterize the expected behavior of the above simulation when we iterate infinite number of times without de-duplication (for the simplicity of the analysis), we developed a program to imitate <u>P</u>roximity <u>L</u>igation by <u>E</u>xpected <u>C</u>onvergence (PLEC), and convert the multi-ChIA complex data into pairwise interactions with adjusting weighting factors generated by PLEC. This approach is based on an assumption that in a Hi-C experiment, all fragments may be ligated during an overnight proximity ligation reaction reaching saturation. Thus, for a complex with *2n* endpoints there would be *n* ligations when saturation is reached. We can further calculate the probability of self-ligations and inter-ligations. Given *n* possible self-ligations, each self-ligation would have a probability 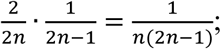 and given 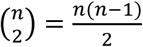 inter-ligations, each inter-ligation would have probability 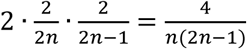. We verify that the total probabilities add up to 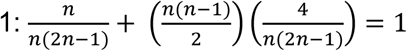. Hence, we can calculate the expected values of the interaction frequencies per complex for all pairs of fragments in the complex by multiplying the individual probabilities by *n*. For self-ligations, the expected values of the interaction frequencies per complex would be 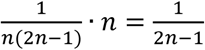. For inter-ligations, the expected values of the interaction frequencies per complex would be 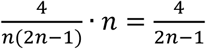. We then use these expected values of the interaction frequencies per complex as weighting factors to convert the multi-ChIA complex data into pairwise Hi-C-like data for fair comparison with previously generated Hi-C data.

Next, we visually compared the adjusted pairwise contacts to the raw pairwise contacts of the multi-ChIA data. We used the following three BEDPE files to generate .hic files for visualization in Juicebox: 1) all pairwise interactions; 2) random sampling with PLISRS weighting factors by 5 iteration ( *k =* 5) and de-duplication; 3) all pairwise interactions adjusted by PLEC weighting factor 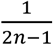 for self-ligations and 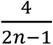 for inter-ligations (where *n* is the number of fragments in the complex). We generated these files for all multiplex GEM complexes with 2 or more fragments (F≥2), with 6 or more fragments (F≥6), and with 10 or more fragments (F≥10) (Extended Data Fig. 3-4). The thresholds 6 and 10 were based on observing two gaps in the empirical cumulative distribution function (ECDF) of interaction distances decomposed by the number of fragments (Fig. 2b).

#### Correlation analysis between multi-ChIA and Hi-C on 2D contact heat maps

We calculated the Pearson correlation coefficient between two heat maps to determine how well the multi-ChIA data correlates with the Hi-C data in regions of interest. For selected chromosome, resolution (25 kb or 5 kb), and normalization method (none or balanced), we extracted contact matrix data from each Juicebox .hic file using ‘dump’ function of Juicer tool. For the normalized (balanced) data, we removed NaN (not a number) values, which were produced by the balanced normalization algorithm for the sparse row and column in the contact matrix. The dumped contact matrix has been already excluded the empty data point in the contact map. Finally, we calculate the Pearson correlation coefficient for the intersection of two dumped contact matrix data sets using C/C++ program.

#### Comparison of pairwise contact spans in various chromatin interaction datasets

To characterize the loop span of multi-ChIA and RNAPII multi-ChIA data, we first randomly sampled 10,000 contacts from all pairwise interactions in each dataset. This set of contacts is further decomposed by the number of fragments in the GEM of its origin. We plotted the genomic spans for empirical cumulative distribution function (ECDF) subgroup of GEMs with 2, 3, 4 fragments (F=2, F=3, F=4), etc. The ECDF curve of Hi-C contacts were included as a reference in the plot for the multi-ChIA data, and the ECDF of RNAPII ChIA-PET contacts was also included as a reference for the RNAPII-enriched multi-ChIA data (Fig. 2b, Extended Data Fig. 9c). The ECDF curves were drawn by ‘stat_ecdf’ function of the ‘ggplot2’ package in R.

#### Identification of RNAPII-associated interaction domains (RAIDs)

Based on chromatin loop intensity and connectivity from the RNAPII ChIA-PET data, we defined RNAPII-associated interaction domains (RAIDs) in *Drosophila* S2 cells in this study. Density plot of RNAPII-associated chromatin interaction loop span (genomic distance) and contact frequency (PET count) shows a bimodal distribution of 2 populations of chromatin contacts, the mid-range from 10 kb to 100 kb, and the long-range larger than 1 Mb (≥1 Mb). The PET count in each chromatin interaction loop reflects the contact frequency. The loops with more PET counts are high frequent and high confidence. Most of the high confidence loops showed high proportion in the mid-range and very low in long-range. On the contrary, most of the low confidence data (with low PET counts) are in long-range. Chromatin loop data with 4 and more PET counts (PET≥4) are considered as high quality for further RNAPII-associated interaction domain (RAID) analysis (Extended Data Fig. 7a). When plotting log10 loop-span distribution, the histogram shows a bimodal Gaussian distribution, which prompted us to fit a mixture of two Gaussian distributions by the Expectation-Maximization algorithm (‘normalmixEM’ function of the R package ‘mixtools’).By observing that the majority of the population belonged to mid-range interactions with mixing coefficients greater than 0.65, we subsequently took 100 kb as a threshold because it is close to the transition point of the mixture models (Extended Data Fig. 7b).This threshold was then used to filter RNAPII loops to be including when demarcating RAIDs. Only those connected domains containing at least three loops were selected as RAIDs structures. Finally, 476 RAIDs were defined (Supplementary Table 3a). Majority of loops with 4 and more PET counts (PET≥4) are located within RAID (intra-RAID loops) with stronger interactions (Extended Data Fig. 7c). There is also a subset of loops that have longer span and connecting RAIDs (inter-RAID loops) and less frequent (Extended Data Fig. 7d).

#### Identification of transcriptional active and inactive domains

To study the relationship between chromatin structure and transcription activity, we defined transcriptionally active and inactive domains. Three data sets in bedGraph format were used as our input: RNA-seq, H3K27ac and H3K27me3 ChIP-seq (Extended Data Table 1). The signal of each data set smoothened using the by applying a rectangular moving averages filter, where the window size is the average gap length (‘filtfilt’ function of the ‘signal’ package in R).These smoothened signals were segmented with the Binary Segmentation algorithm (‘cpt.meanvar’ function with Q = 100, minseglen = 5000 of the ‘changepoint’ package in R).We retained the segments with estimated mean greater than 5 and recorded its chromosome name, start position, end position, and estimated mean. The three resulting bedGraph files were concatenated (‘unionbedg’ function with –empty of the bedtools^24^), where each line has a set of unique estimated mean from RNA-seq, H3K27ac, and H3K27me3, along with its chromosome name, start position, and end position.Intuitively, a segment with high RNA-seq and H3K27ac signal coupled with low H3K27me3 signal implies a transcriptionally active region.Next, we iteratively merged small nearby regions to obtain 708 active regions across chromosomes 2L, 2R, 3L, 3R, 4, and X.Finally, the complements of active regions are defined as inactive regions (‘complement’ function of the bedtools^24^) (Supplementary Table 3b-c). An example is shown in Figure 3a.

#### Chromatin fragments in multi-ChIA data related to TADs

To relate the multi-ChIA data to TADs defined by Hi-C data^18^, multiplex chromatin complexes with fragments 6 or more (F≥6) were hit to TADs, and at least 3 chromatin fragments of a complex located in a TAD would be regarded as enriched in the TAD region. If an individual chromatin complex contained fragments overlapped with 2 or more TADs (at least 3 fragments in each TAD), it would be considered as crossing TADs. In the complexes (F≥6), 85% of them were enriched in 1 TAD, 11% of them were crossed 2 TADs and 4% were crossed multiple TADs. This result is presented in Figure 2e.

#### Chromatin fragments in RNAPII multi-ChIA data related to RAIDs

The chromatin complexes (GEMs) with 3 or more fragments (F≥3) detected by RNAPII multi-ChIA were hit to RAIDs defined by RNAPII ChIA-PET data (Supplementary Table 3a). Three categories were identified, GEMs overlap 0 RAIDs indicated no any fragments of individual GEM located in RAIDs, GEMs overlap 1 RAIDs indicated there was one fragment located in RAIDs, GEMs overlap ≥ 2 indicated more than one fragments located in RAIDs. The result is presented in Extended Data Figure 11.

#### Comparison of multi-ChIA and RNAPII multi-ChIA data

The chromatin fragments in multi-ChIA and RNAPII multi-ChIA data were hit to TADs^18^ and RAIDs, respectively (Supplementary Table 3a). Total fragment numbers hit in the two types of domains were used to calculate the normalized density for FPKM (fragment per kb and per million reads) in boxplots. The result is presented in Figure 3c. Similarly, the chromatin fragments were also hit to transcriptionally active and inactive domains, respectively, and calculated for FPKM in boxplot. The distributions of FPKM values were visualized using box plots (Extended Data Fig. 10). The Wilcoxon Rank-Sum Test was used to assess the significance of differences.

#### Enhancers and promoters in RNAPII multi-ChIA data

RNAPII multi-ChIA detected multiplex chromatin complexes with 3 or more fragment (F≥3) and overlapped with RAIDs were selected for analysis. Fragment overlapped with gene TSS region (± 250 bp) was assigned as promoter^28^, fragment overlapped with TSS distal region was considered as potential enhancer. In the defined 11,695 complexes, there are 40% belonging to 2 enhancers with 1 promoter (abbreviated as 2E1P), 34% was involved in multiple enhancers with 1 promoter (mE1P), and 9% belonging to 1 enhancer with 2 promoters (1E2P). The rest of them include multiple promoters with multiple enhancers (Fig. 3e; Supplementary Table 4).

#### Data availability

The multi-ChIA and RNAPII multi-ChIA data (raw sequencing files FASTQ and BAM, processed data files of uniquely mapped reads with GEMcodes, and files of chromatin interaction complexes) and ChIA-PET data (raw sequencing files FASTQ and BAM, files of reads pileup coverage, files of interaction clusters) are available at Gene Expression Omnibus under SuperSeries accession number: GSEXXXX. And publically available datasets used in this study are reported in Extended Data Table 1.

## Extended Data Information

12 Extended Data Figures

1 Extended Data Table

**Extended Data Figure 1.**
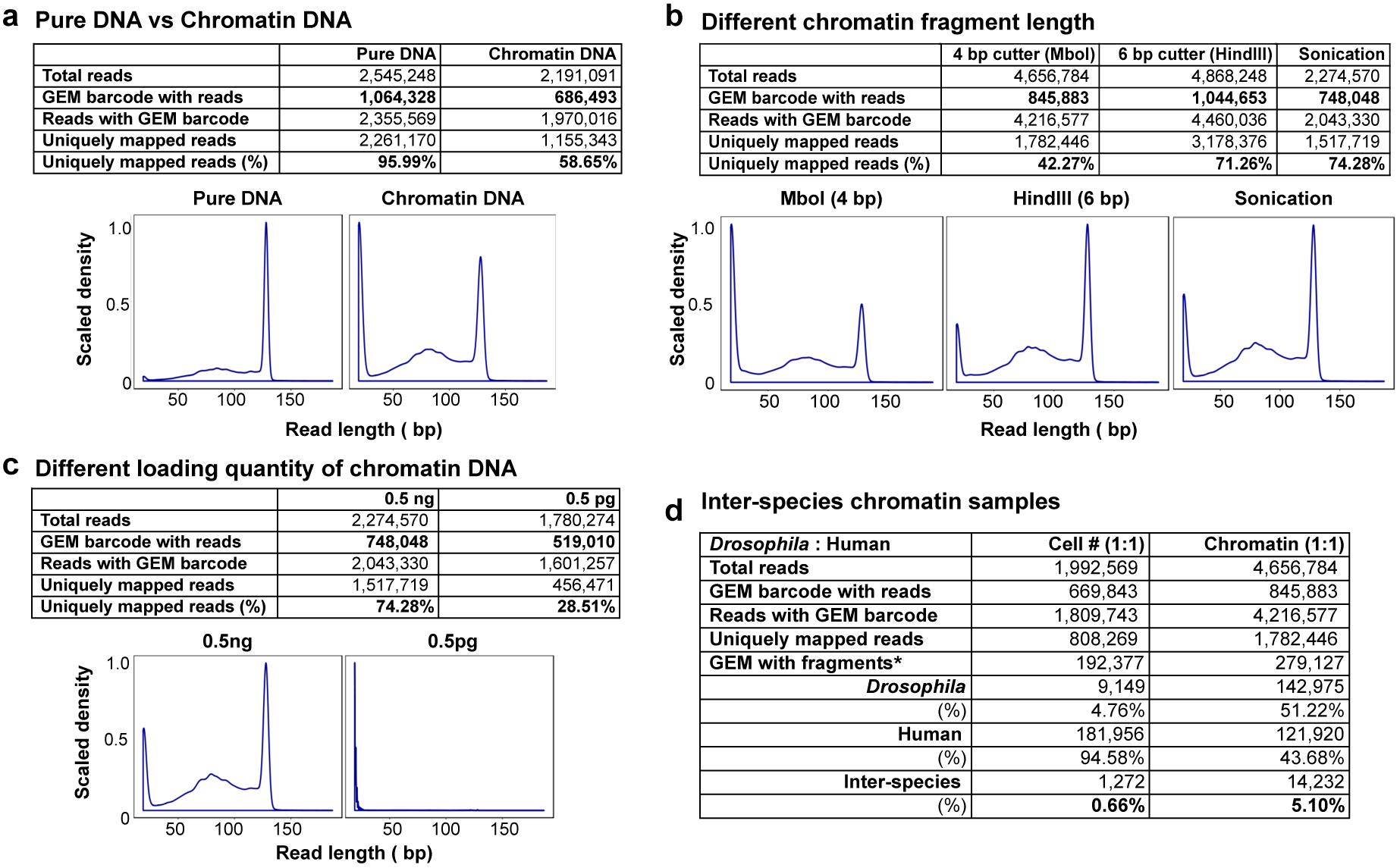
Characterization of Chromium microfluidics system for chromatin DNA sequencing. The efficiency of the microfluidics system for chromatin DNA barcoding and amplification was characterized by MiSeq sequencing data. Each test generated 2-4 million sequencing reads. The numbers of captured GEM barcodes and the percentages of uniquely mapped reads (in tables) and the read length distribution (in plots) are indicative for data quality. **a**, Purified DNA versus chromatin DNA. Both DNA templates were prepared from the same chromatin sample. Using pure DNA templates as a performance reference (note: this 10x Chromium microfluidics system is optimized for genomic DNA molecules), 58.65% of the reads generated from chromatin fragments are useful for chromatin analysis. **b**, Different chromatin fragment length. Chromatin sample fragmented by 4 bp cutter (MboI, ∼300 bp), and 6 bp cutter (HindIII, ∼3000 bp), or sonication (∼6000 bp) were prepared accordingly. The results indicate that longer chromatin fragments are better suited for chromatin analysis in this microfluidics system than the shorter ones. **c**, Chromatin sample loading with different input quantity, 0.5 ng versus 0.5 pg. If the input is too low (i.e., 0.5 pg), most of the sequencing reads were only 19-20 bp, indicating that most of the droplets were empty of chromatin materials. **d**, Inter-species chromatin samples. Chromatin samples from *Drosophila* S2 and human GM12878 cells were mixed in equal cell numbers or in equal chromatin mass. Barcoded sequencing reads mapped to each reference genome were merged into fragment based on overlapping alignment in each GEM. The numbers of GEMs with fragments of genomic origins (*Drosophila*, human, or mixed) were counted. As noticed, in the test with equally mixed numbers of cells, the number of GEMs with chromatin fragments of human origin is 20-fold more than GEMs of *Drosophila* origin (181,956 / 9,149 = 19.89), which is in approximate match with the genome size difference between human and *Drosophila* (hg 3,000 Mb / dm 175 Mb = 17.14). In the test with equally mixed chromatin mass, the numbers of GEMs of human origin and of *Drosophila* origin are close to 1:1 ratio. Notably, the GEMs with mixed origins of fragments were only 5.1%, indicating a small rate of individual droplets that may contain mixed chromatin samples.

**Extended Data Figure 2.**
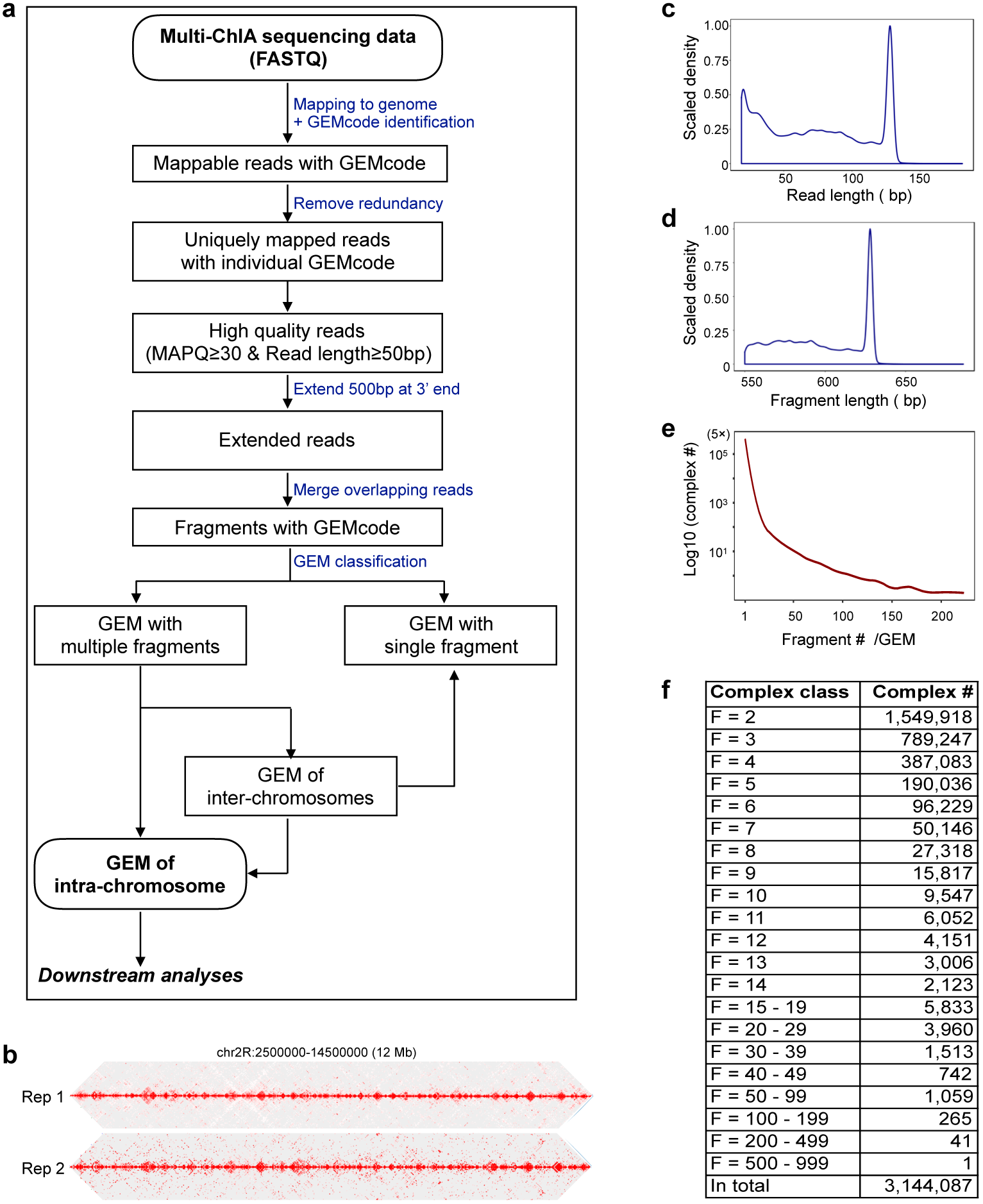
Multi-ChIA data processing pipeline and sequencing profile. **a**, Flowchart of multi-ChIA library sequencing data processing pipeline. **b**, 2D chromatin contact views for two multi-ChIA replicates (Rep1 and Rep2). The reproducibility of the two replicates was assessed by Pearson correlation coefficient analysis (*r* = 0.75). **c**, Size distribution plot of sequencing read length. Only R1 reads that contain GEMcodes and genomic sequences were used for analysis in this study. The maximum read length is 130 bp plus related linker information. The shortest reads were in 20 bp, most of which were residuals from linker dimers. **d**, Size distribution plot of merged chromatin fragment length. The peak length is approximate 630 bp. **e**, The distribution plot of numbers of merged chromatin fragments in individual GEMs. **f**, GEMs contain multiple chromatin fragments (F≥2) are considered as multiplex GEMs, inferred as chromatin complexes. Multiplex complexes were categorized based on the numbers of fragments in each complex. F=2, chromatin complexes with 2 fragments; F=3, chromatin complexes with 3 fragments; and so forth.

**Extended Data Figure 3.**
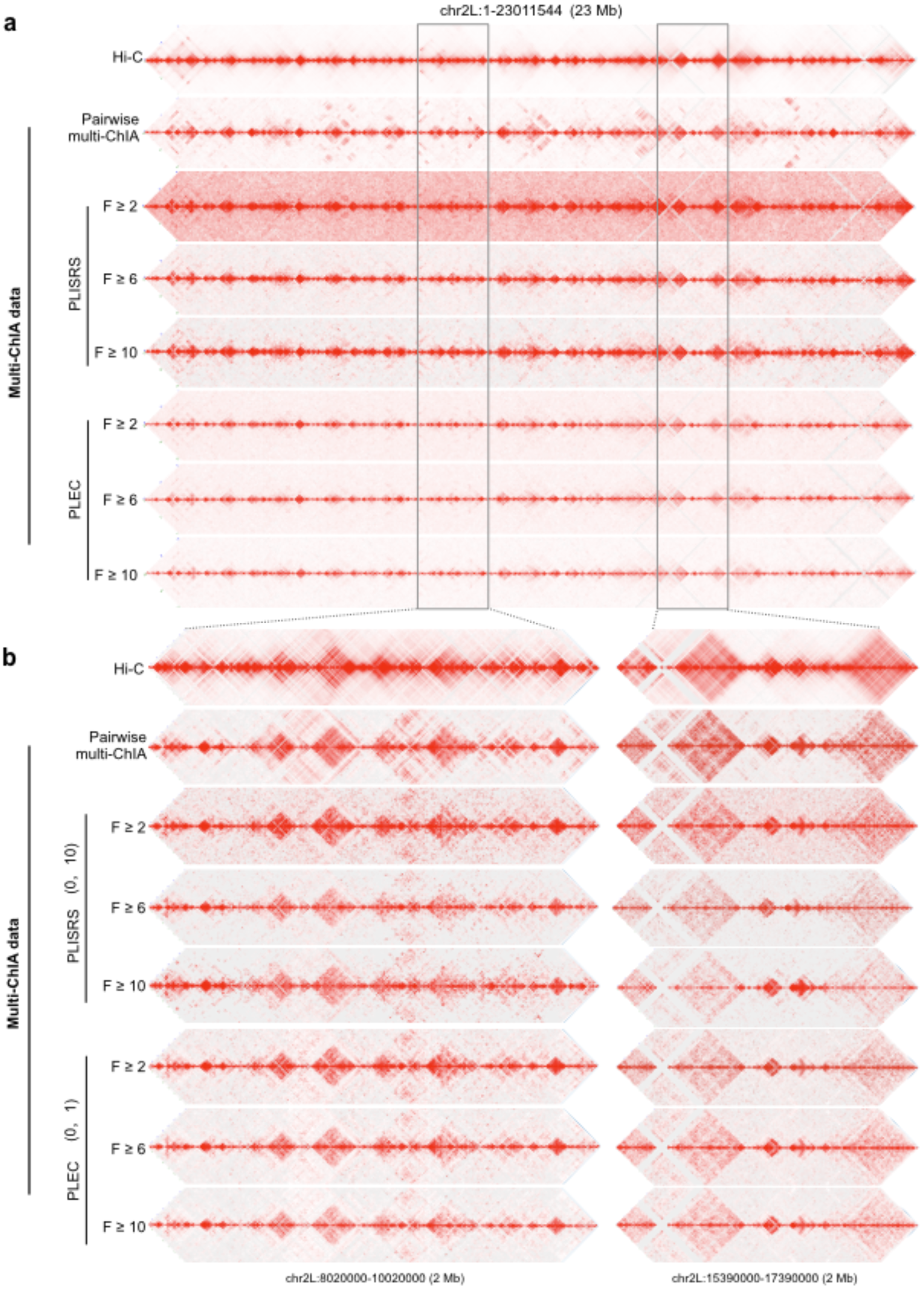
Adjustment of multi-ChIA pairwise data by weighting factors for comparison with Hi-C data. **a**, 2D contact view of chromosome 2L at 25 kb bin-size resolution of Hi-C and multi-ChIA data. Pairwise-based multi-ChIA data were adjusted with various weighting factors: Pairwise multi-ChIA data (raw contacts, no adjusted weighting factors applied); PLISRS (<u>P</u>roximity <u>L</u>igation *<u>I</u>n <u>S</u>ilico* by <u>R</u>andom <u>S</u>ampling) weighting factors adjusted multi-ChIA complex data with different fragment classes (F ≥ 2, F ≥ 6 and F ≥10), respectively; and PLEC (<u>P</u>roximity <u>L</u>igation by <u>E</u>xpected <u>C</u>onvergence) weighting factors adjusted multi-ChIA complex data of same fragment classes. The chromatin contact overall profiles between the Hi-C data and the multi-ChIA data are significantly in consistence (Pearson correlation coefficient *r* > 0.7). **b**, Zoom-in views on a 2 Mb segment at 5 kb bin-size resolution.

**Extended Data Figure 4.**
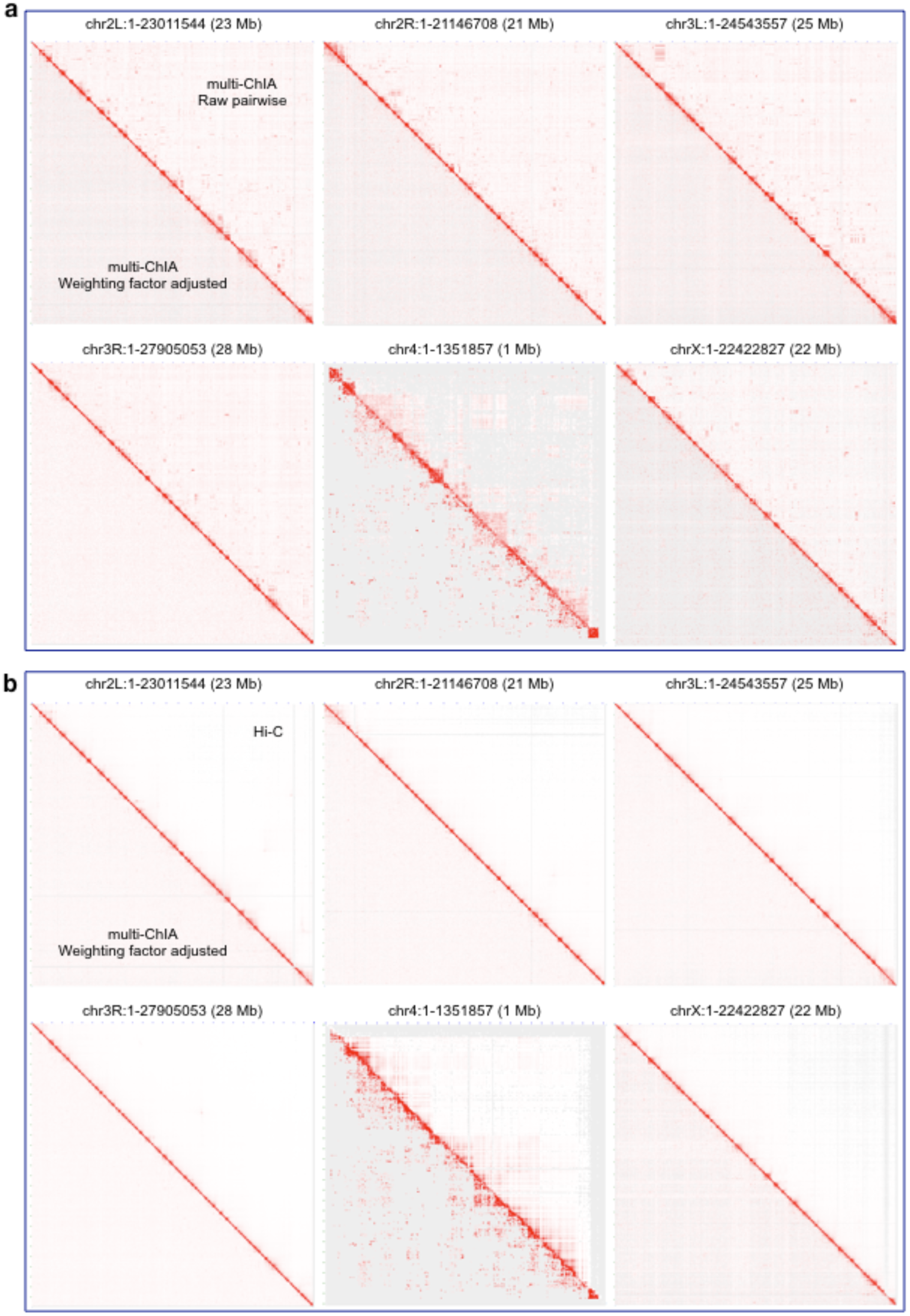
Genome-wide comparison of multi-ChIA data with Hi-C data. a, 2D contact views of multi-ChIA pairwise data before (Raw pairwise) and after adjustment (Weighting factors adjusted) for the 6 chromosomes in Drosophila genome. The “Raw pairwise” 2D contact views are at upper diagonal, and the “Weighting factor adjusted” 2D contact views are at lower diagonal. It is evident that the adjustment by weighting factors calculated by the PLEC algorithm could help to smooth the 2D contact views of multi-ChIA data. b, Comparison of the Hi-C data (upper diagonal) and the weighting factor adjusted multi-ChIA pairwise data. The multi-ChIA data in pairwise contacts were adjusted based on PLEC weighting factors (see Methods) using data of chromatin complexes with six or more fragments (F≥ 6) at 25 kb bin-size resolution (Pearson correlation coefficient r > 0.8).

**Extended Data Figure 5.**
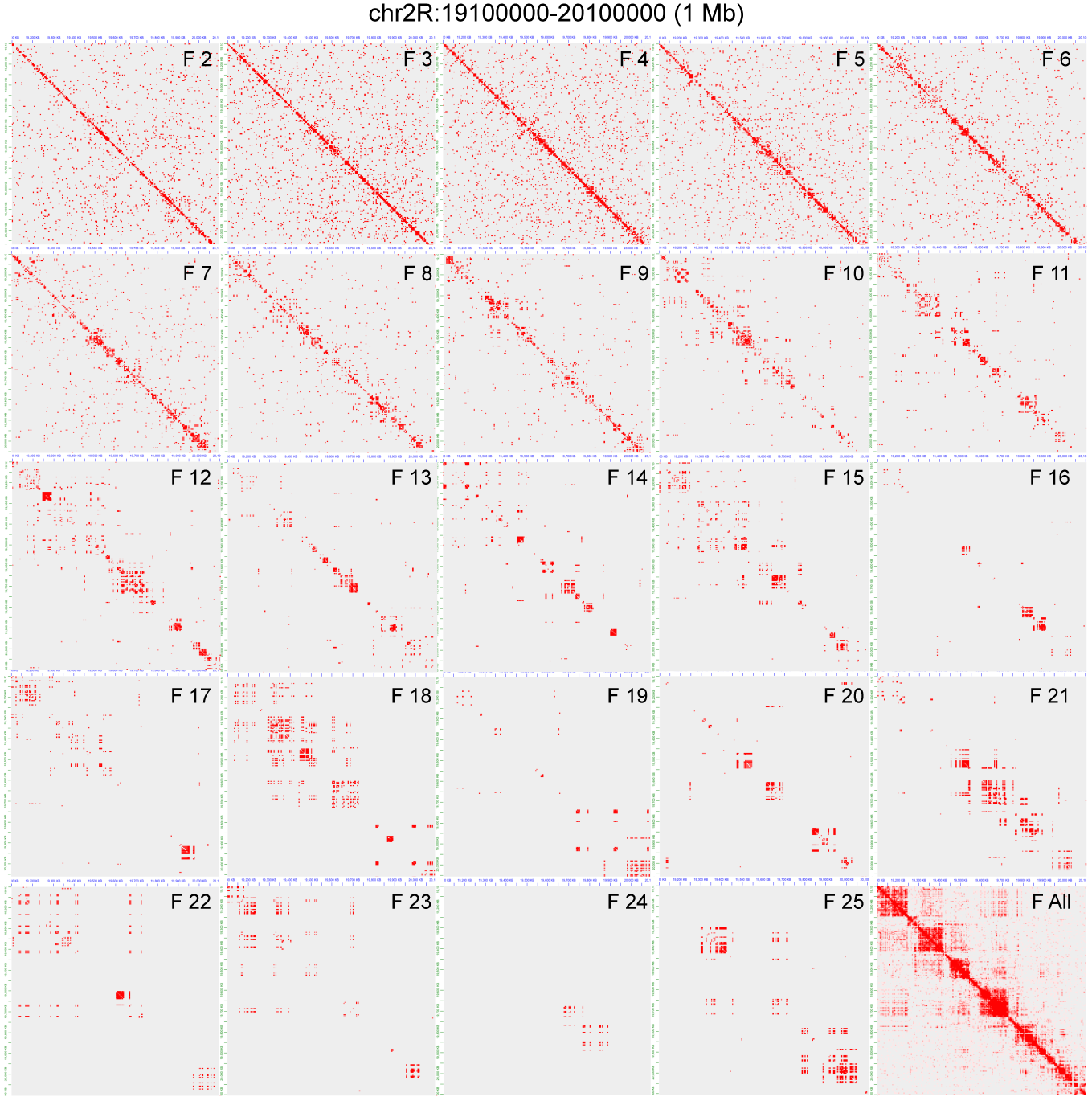
Decomposition analysis of multi-ChIA data. Shown are 2D contact views of decomposition of multi-ChIA data based on the number of DNA fragments in each chromatin complex. F2 (complex data with 2 fragments), F3 (complex data with 3 fragments), and so forth, up to F25 (complex data with 25 fragments), and F All for all complex data as an overall reference.

**Extended Data Figure 6.**
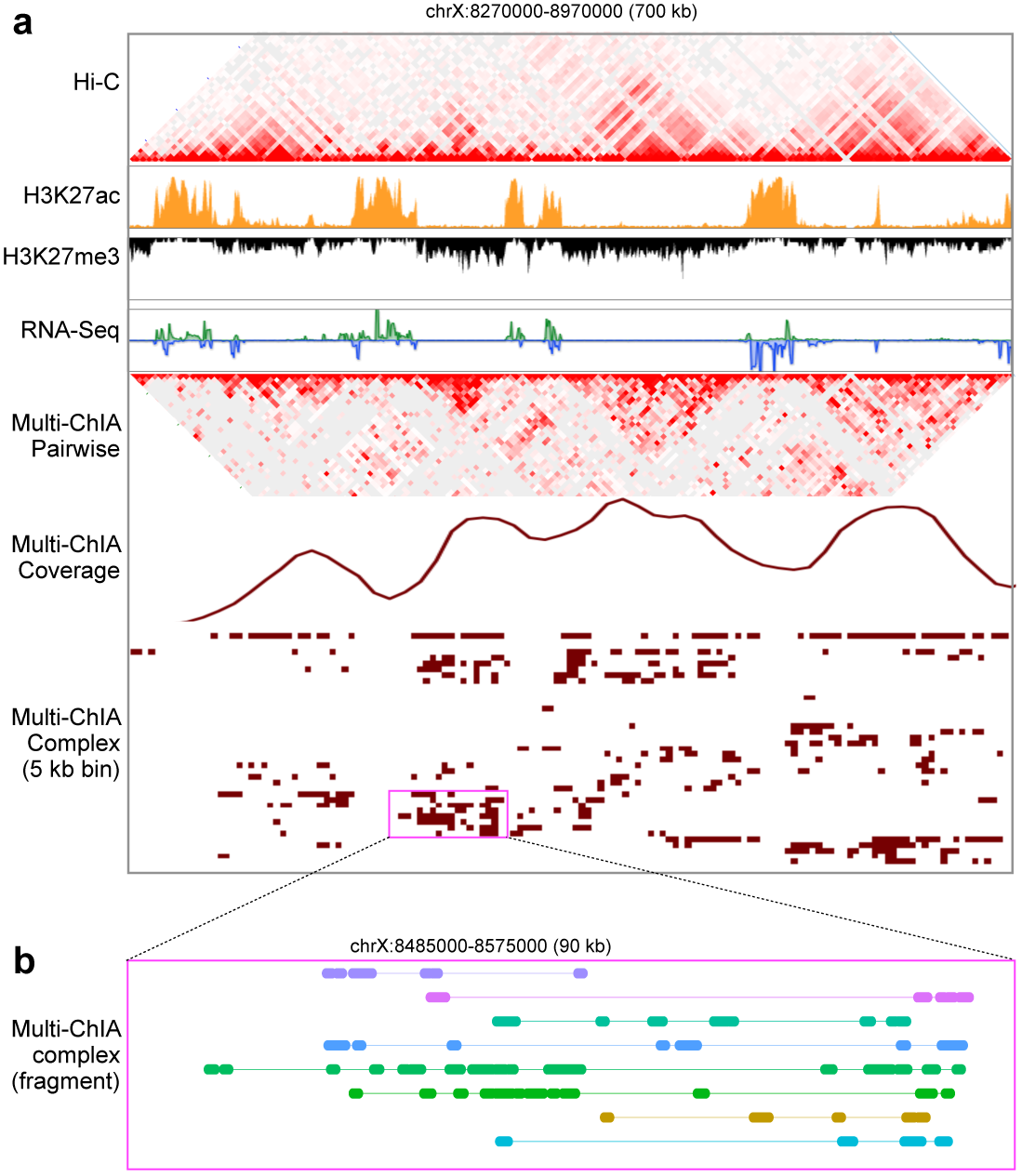
The multi-ChIA data visualization. **a**, An example view of the multi-ChIA data at a 700 kb segment in chromosome X of *Drosophila.* The Hi-C data outlined 4 TAD structures in this region, along with histone and transcriptional profiles to reference the chromatin states. The multi-ChIA data of pairwise contacts showed comparable contact structures resembling the Hi-C data. A density curve plotted the overall multi-ChIA data coverage in this region. The chromatin fragments in multiplex complexes were linearly aligned across this segment with 5 kb bin-size resolution, and the hierarchical-clustering displayed enrichment of multi-ChIA data in TAD domains. **b**, A zoom-in of 90 kb region displays detailed linear alignment of 8 chromatin complexes with multiple fragments in assorted color bars.

**Extended Data Figure 7.**
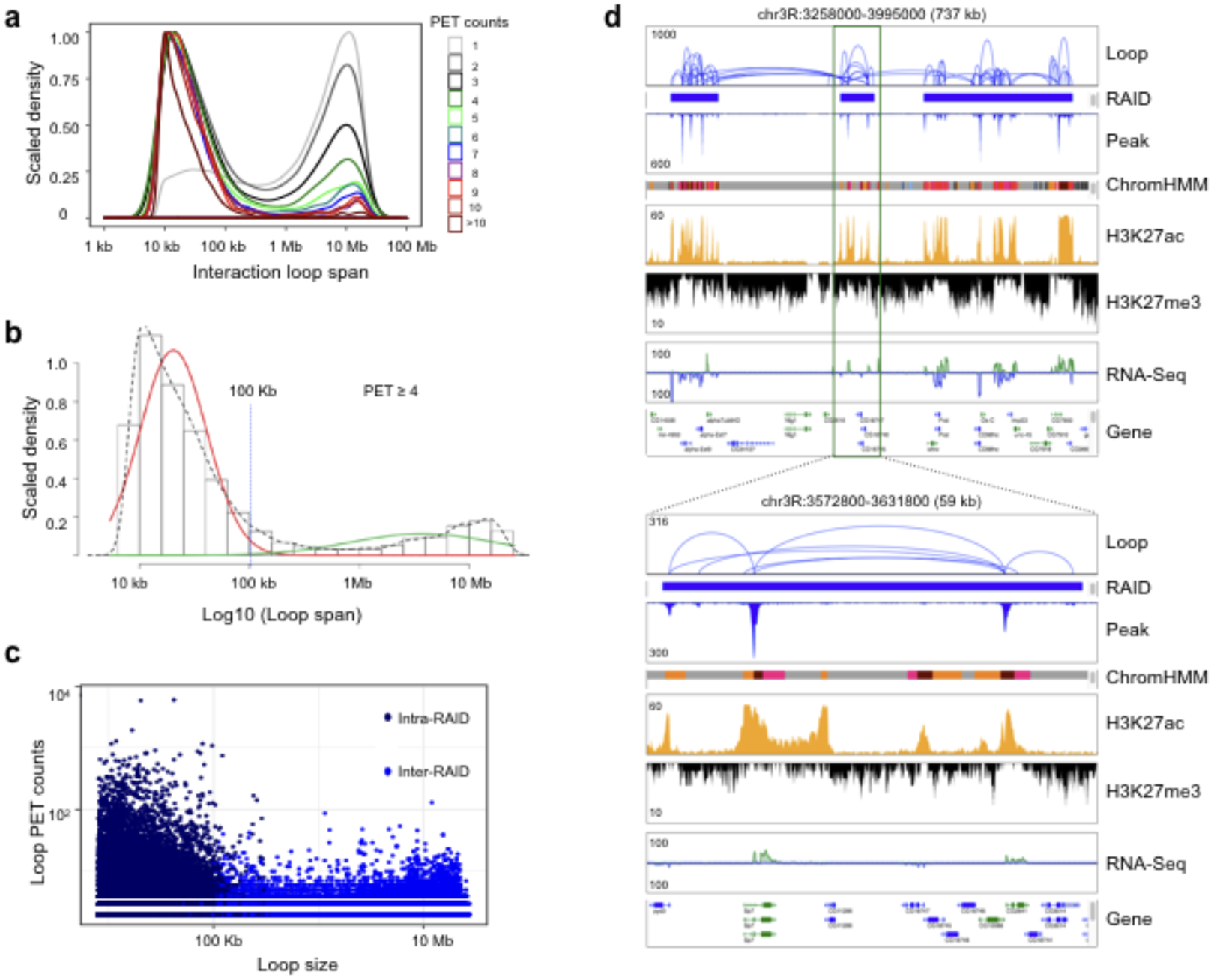
Definition and characterization of RNAPII-associated interaction domain (RAID). **a**, Density plot of RNAPII-associated chromatin interaction loop span (genomic distance) and contact frequency (PET count) shows a bimodal distribution of 2 populations of RNAPII loops based on loop span, the mid-range from 10 kb to 100 kb, and the long-range larger than 1 Mb (≥1 Mb). The PET count in each RNAPII loop reflects the contact frequency. PET 1 (light grey) stands for a contact data with 1 PET (PET singleton), PET2 (grey) for contact with 2 PETs, PET3 (black) with 3 PETs, and so forth. The loops with more PET counts are high confidence and reflect high frequency contacts. Most of the high confidence loops showed high proportion in the mid-range and low in long-range. On the contrary, most of the low confidence data (with less PET counts) are in long-range. Chromatin loop data with 4 and more PET counts (PET≥4) are considered as high quality for further RNAPII-associated interaction domain (RAID) analysis. **b**, Histogram of statistical analysis on RNAPII associated chromatin loop span (in log scale) with 4 or more PET counts (PET≥4). The population of loops is separated into the mid-range interactions (red) and the long-range interactions (green) through the Gaussian mixture model. Dotted line shows the marginal distribution, and 100 kb loop span was taken as a threshold to separate the two populations. **c**, Scatter plot of loop span and contact frequency of the chromatin interactions that located within RAIDs (intra-RAID, dark blue) and the chromatin interactions that connecting RAIDs (inter-RAID, light blue). Most intra-RAID loops were stronger interactions (higher PET counts) and within 100 kb, whereas the inter-RAID loops were mostly weaker (lower PET counts) and longer span. **d**, An example of browser view shows the defined RAIDs, RNAPII loops within RAIDs and connecting RAIDs (blue). Tracks of chromatin state (ChromHMM: red for promoter; dark for enhancer; grey for inactive region), histone marks (ChIP-Seq for H3K27ac and H3K27me3), and transcription activity (RNA-seq) are included for epigenomic reference. Gene model track also included for genomic reference. A zoom-in view highlights the details of RNAPII associated chromatin loops within a RAID.

**Extended Data Figure 8.**
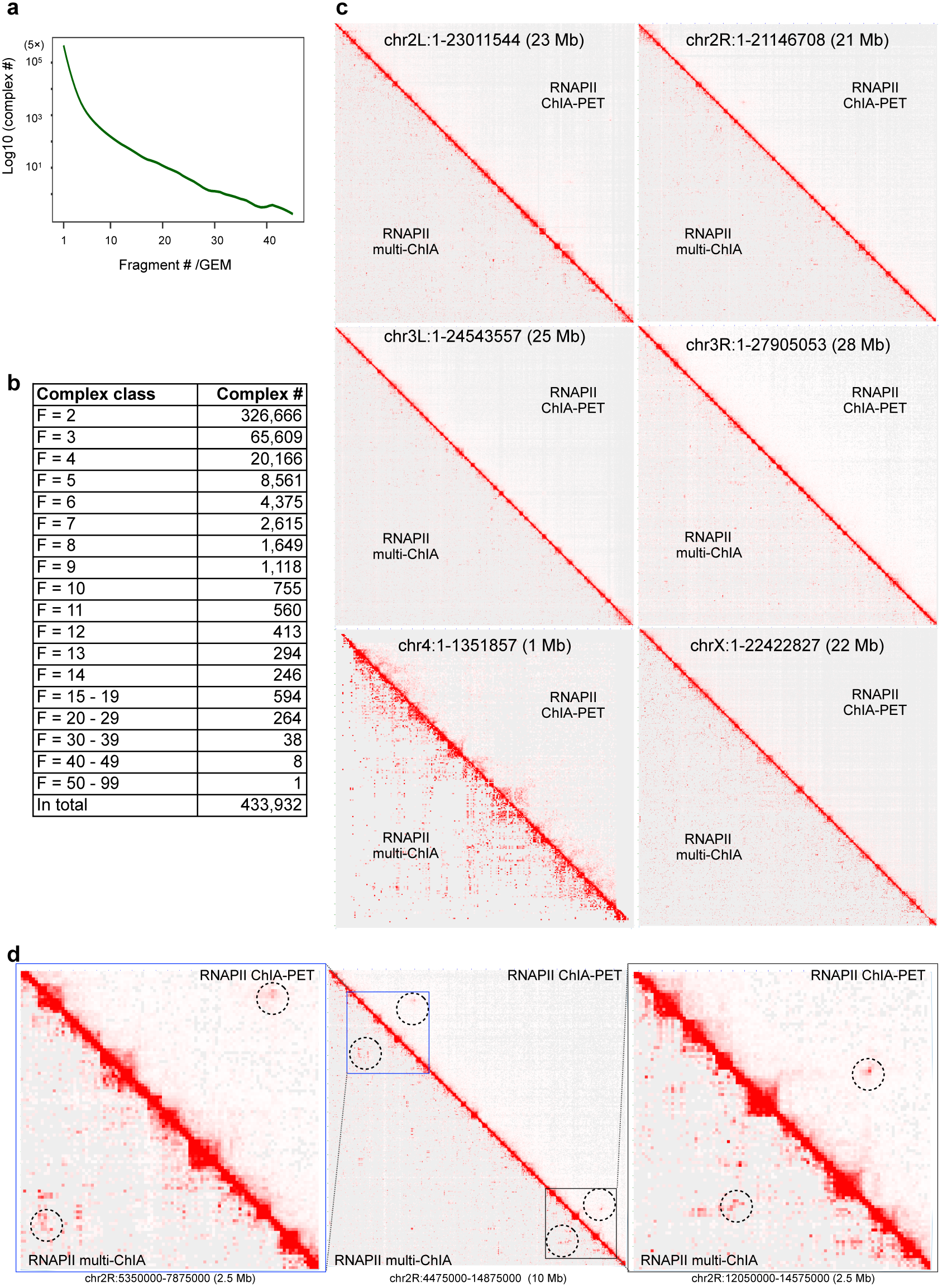
Characterization of RNAPII multi-ChIA data. **a**, Distribution plot of chromatin fragment numbers in individual GEMs detected by RNAPII multi-ChIA. **b**, Multiplex chromatin complexes with various fragment numbers. F=2, chromatin complexes with 2 fragments; F=3, chromatin complexes with 3 fragments; and so forth. **c**, 2D contact views of the six chromosomes in *Drosophila* genome between RNAPII ChIA-PET data (upper diagonal) and the adjusted pairwise contacts of RNAPII multi-ChIA data (lower diagonal) with 25 kb bin-size resolution (Pearson correlation coefficient *r* > 0.75). **d**, Zoom-in views of 3 different segments in chromosome 2R (chr2R) with 25 kb bin-size resolution. Corresponding circles highlight specific chromatin contacts in both of the RNAPII ChIA-PET data and the RNAPII multi-ChIA data (Pearson correlation coefficient *r* > 0.78).

**Extended Data Figure 9.**
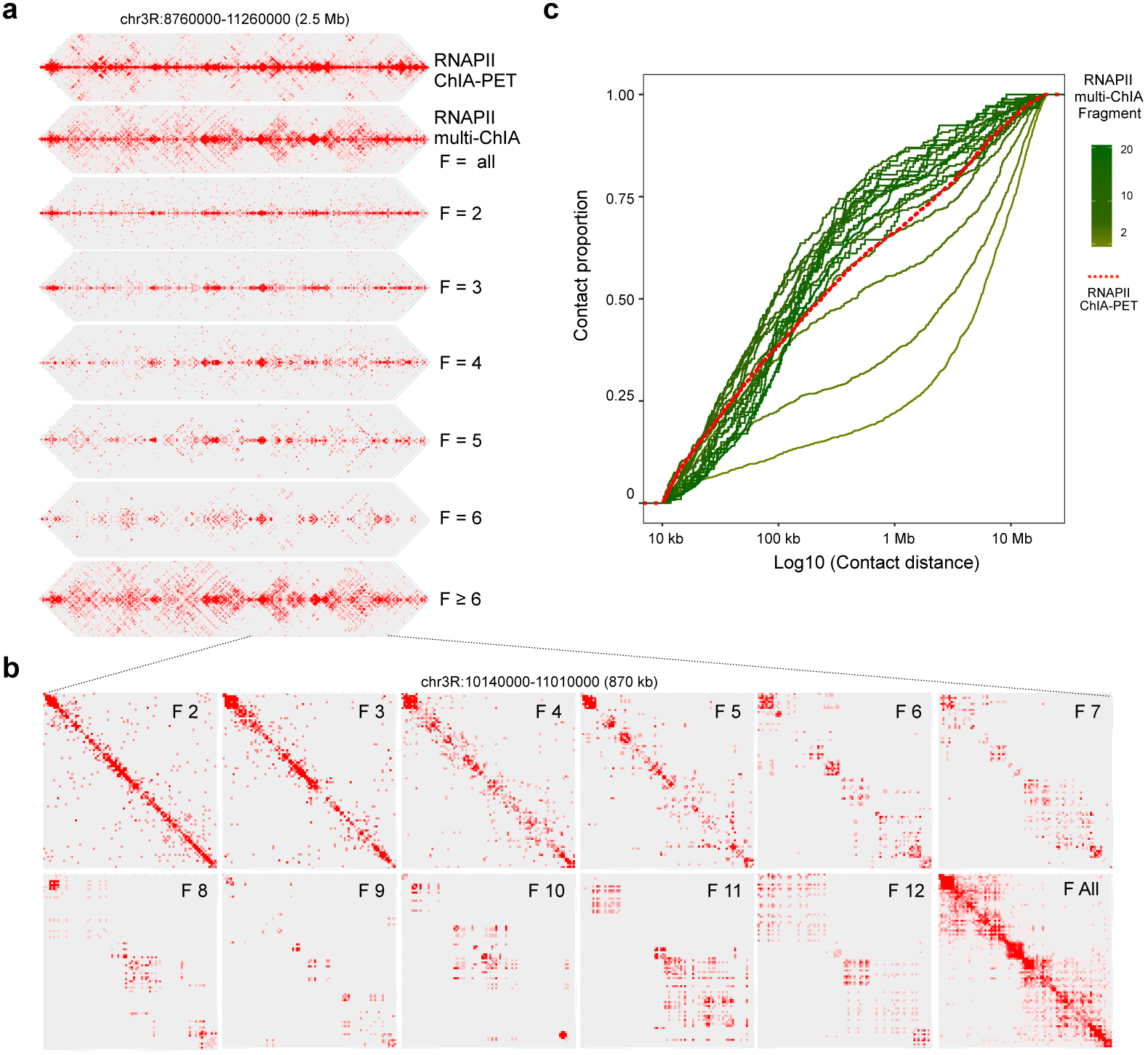
Decomposition analysis of RNAPII multi-ChIA data. **a**, Decomposition of RNAPII multi-ChIA data based on the number of fragments in each chromatin complex. F=all, all complex data; F=2, complexes with 2 fragments; F=3, complexes with 3 fragments; F≥6, complexes with 6 or more fragments. The 2D contact views are for a 2.5 Mb segment in chromosome 3R (chr3R) with 5 kb bin-size resolution. RNAPII ChIA-PET data is included as a reference. **b**, Zooming-in 2D contact views from (**a**) of an 870 kb segment with 5 kb bin-size resolution for the decomposed RNAPII multi-ChIA data based on the number of fragments in each chromatin complex. **c**, Empirical cumulative distribution function (ECDF) of chromatin contact proportion versus chromatin contact distance in log scale. The solid green curves denote RNAPII multi-ChIA data of pairwise contacts with increased color intensity corresponding to different numbers of fragments per complex from low to high. The dotted red curve depicts *in situ* RNAPII ChIA-PET data of pairwise contacts as a reference.

**Extended Data Figure 10.**
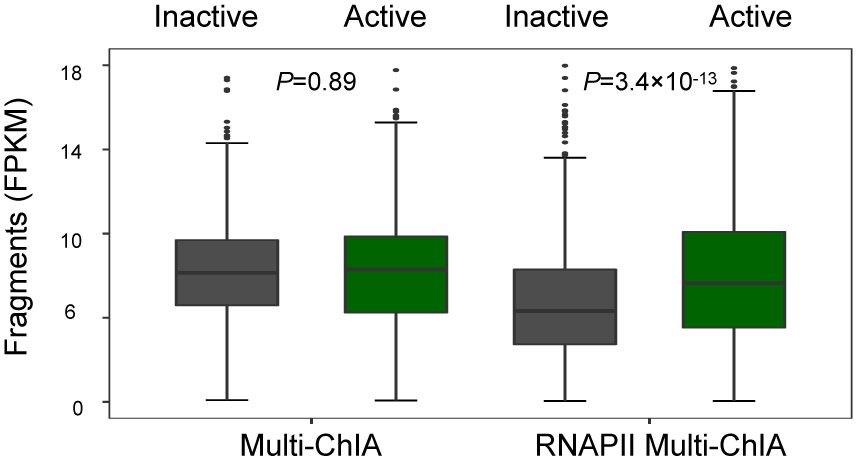
Multi-ChIA data related to active and inactive domains. Boxplots of normalized chromatin fragments (FPKM) in multi-ChIA and RNAPII multi-ChIA data represented in the defined active or inactive domains. In multi-ChIA data, the normalized chromatin fragments (FPKM) showed no difference. As for a comparison, the normalized fragments in the RNArll multi-ChIA data were significantly *(p=3.4x10)* reduced in inactive domains and increased in active domains. Wilcoxon Rank-Sum Test was used to test the significance.

**Extended Data Figure 11.**
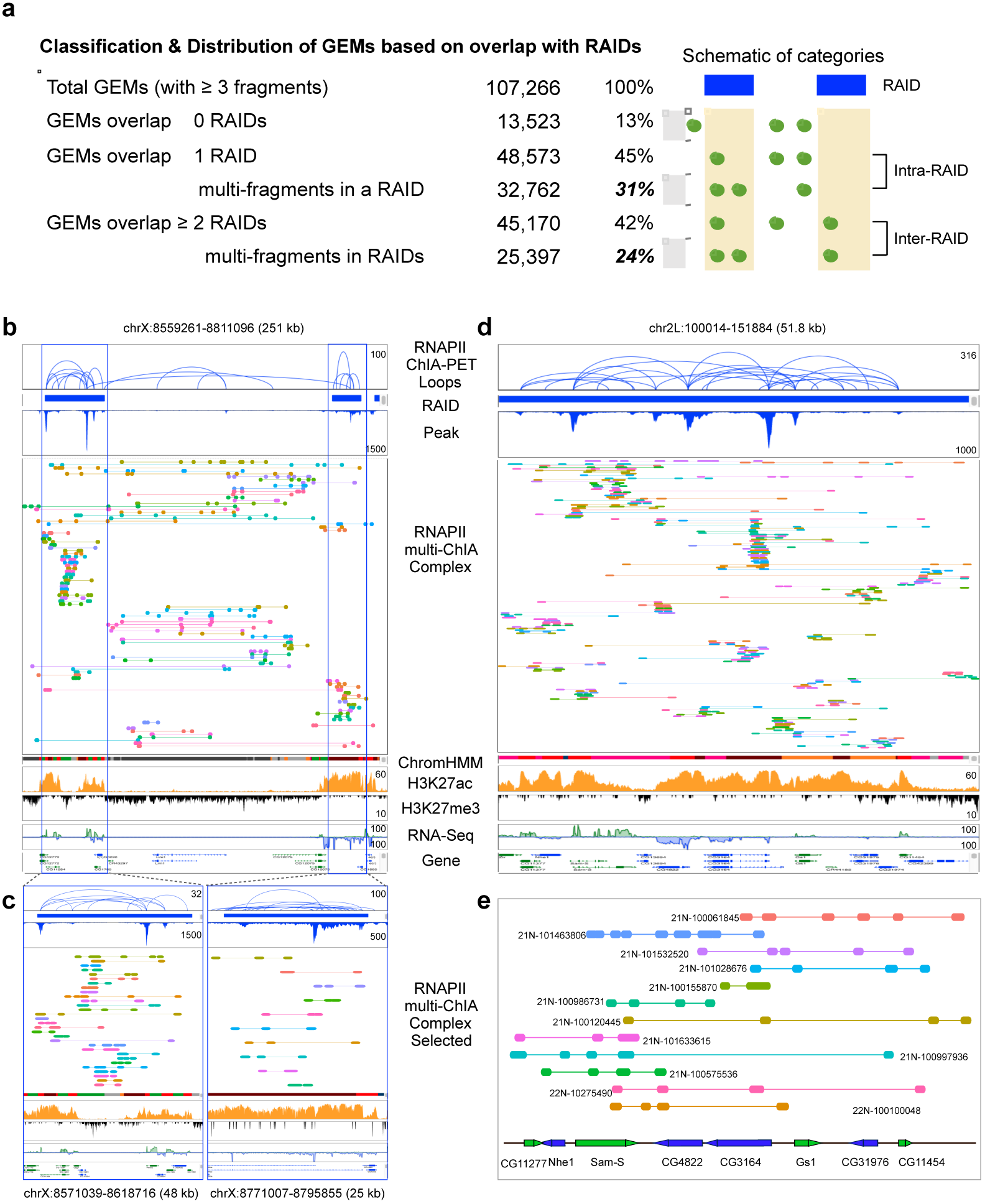
Chromatin complexes overlapping with RAIDs. **a**, Considering all chromatin complexes (GEMs) with 3 or more fragments (F≥3), 87% of the complexes are overlapped with RAIDs, and 42% of them overlapped with 2 or more RAIDs. Blue bars denote RAIDs, green dots denote chromatin fragments, and the green lines connecting the dots depict chromatin complexes (GEMs). **b**, Screenshot of a browser view in a 251 kb segment in chrX showing RNAPII ChIA-PET data identified RNAPII loops within 2 RAIDs and connecting the 2 RAIDs, along with multi-ChIA data showing the connected chromatin fragments in individual multiplex chromatin complexes across this segment. Tracks of chromatin state (ChromHMM: red for promoter; dark for enhancer; grey for inactive region), histone marks (ChIP-Seq for H3K27ac and H3K27me3), and transcription activity (RNA-seq) are included to exhibit the active and inactive chromatin states in association with RAID and inter-RAID regions. Gene model track is also included for genomic reference. **c**, Zooming-in views of (**b**) on the 2 RAIDs showing further detailed alignment of chromatin fragments in multiplex complexes. **d**, Another example showing RNAPII loops within a RAID and the linear alignment of chromatin fragments of multiplex complexes detected by RNAPII multi-ChIA data. There are 7 active genes reside in this region in S 2 cells. **e**, Selected individual chromatin complexes (with specific GEM IDs) from (**d**) are highlighted for their fragments in relation to gene positions.

**Extended Data Figure 12.**
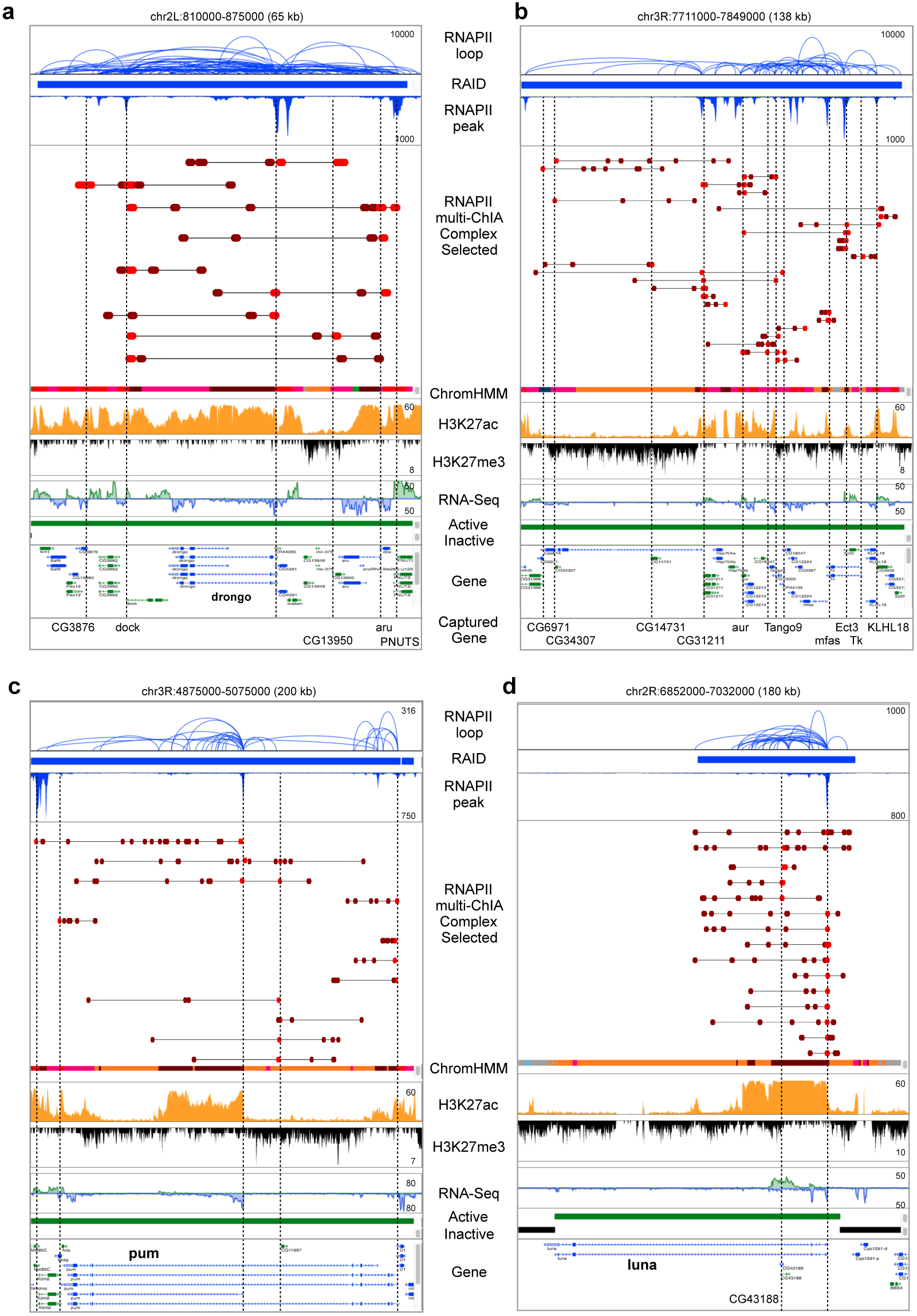
Alignment of RNAPII multi-ChIA complex data to enhancers and promoters in RAIDs. **a**-**d**, Additional examples of multiplex chromatin interactions mapped within RAIDs. Red-colored bars denoted for chromatin fragments overlapped with TSS sites as promoter (P), and the brown-colored bars for chromatin fragments overlapped with TSS distal regions as potential enhancer elements. The TSS sites of genes are highlighted by dotted vertical lines.

**Extended Data Table 1.** Summary of datasets used in this study.

| Method | Library ID | Total reads | Uniquely mapped reads | Data source |
| --- | --- | --- | --- | --- |
| Multi-ChIA | Multi-ChIA_1 | 71,667,907 | 65,810,896 | This study |
| Multi-ChIA | Multi-ChIA_2 | 71,901,950 | 66,042,256 |  |
| Multi-ChIA | Multi-ChIA_3 | 71,182,692 | 65,177,026 |  |
| Multi-ChIA | Multi-ChIA_4 | 71,202,104 | 65,070,408 |  |
| RNAPII_Multi-ChIA | RNAPII-multi-ChIA_1 | 44,728,172 | 16,887,291 |  |
| RNAPII_Multi-ChIA | RNAPII-multi-ChIA_2 | 46,991,737 | 22,471,185 |  |
| RNAPII-ChIA-PET | RNAPII-ChIA-PET_1 | 18,539,755 | 12,370,083 | This study |
| RNAPII-ChIA-PET | RNAPII-ChIA-PET_2 | 20,573,763 | 13,053,275 |  |
| *In situ* RNAPII-ChIA-PET | In situ-RNAPII-ChIA-PET_1 | 15,125,381 | 10,741,175 |  |
| *In situ* RNAPII-ChIA-PET | In situ-RNAPII-ChIA-PET_2 | 10,014,646 | 7,320,147 |  |
| Hi-C | GSM1690083; GSM1690084 |  |  | Ulianov et al., 2016 |
| Hi-C | GSM1420136; GSM1420137 |  |  | Ramirez et al., 2015 |
| ChIP-seq (H3K27ac) | GSM1017404 |  |  | Herz et al., 2012 |
| ChIP-seq (H3K27me3) | GSM480157 |  |  | Gan et al., 2010 |
| Chromatin states |  |  |  | modEncode<br>Kharchenko et al.2011 |
| RNA-seq | SRR1197280 |  |  | Duff et al., 2015 |
| RNA-seq | GSM1000412 |  |  | Arnold et al., 2013 |

## Supplementary Information 4

### 4 Supplementary Tables

Supplementary Table 1 **Multi-ChIA detected chromatin complexes (F ≥ 10).** Related to Extended Data Figure 2f.

Supplementary Table 2 **RNAPII multi-ChIA detected chromatin complexes (F ≥ 6).** Related to Extended Data Figure 8b.

Supplementary Table 3 **Summary statistics of chromatin domains**

Supplementary Table 3a **Summary statistics of RNAPII associated interaction domains (RAIDs).** Related to Figure 3a and Extended Data Figure 7.

Supplementary Table 3b **Summary statistics of Active transcribed domain.** Related to Figure 3a and Extended Data Figure 10.

Supplementary Table 3c **Summary statistics of Inactive transcribed domain.** Related to Figure 3a and Extended Data Figure 10.

Supplementary Table 4 **RNAPII multi-ChIA detected enhancers and promoters involved in multiplex chromatin interactions.** Related to Figure 3e.

